# Unique growth and morphology properties of Clade 5 *Clostridioides difficile* strains revealed by single-cell time-lapse microscopy

**DOI:** 10.1101/2024.02.13.580212

**Authors:** John W. Ribis, César Nieto, Nicholas V. DiBenedetto, Anchal Mehra, Pola Kuhn, Qiwen Dong, Irene Nagawa, Imane El Meouche, Bree B. Aldridge, Mary J. Dunlop, Rita Tamayo, Abhyudai Singh, Aimee Shen

## Abstract

*Clostridioides difficile* is a gastrointestinal pathogen of both humans and agricultural animals and thus a major One Health threat. The *C. difficile* species consists of five main clades, with Clade 5 currently undergoing speciation from Clades 1-4. Clade 5 strains are highly prevalent in agricultural animals and can cause zoonotic infections, suggesting that these strains have evolved phenotypes that distinguish them from Clade 1-4 strains. Here, we compare the growth properties of Clade 5 strains to those of Clade 1-4 strains using anaerobic time-lapse microscopy coupled with automated image analysis. Our analyses indicate that Clade 5 strains grow faster and are more likely to form long chains of cells than Clade 1-4 strains. Using comparative genomic and CRISPRi analyses, we show that the chaining phenotype of Clade 5 strains is driven by the orientation of the invertible *cmr* switch sequence, with chaining strains exhibiting a bias to the *cmr-*ON state. Interestingly, Clade 5 strains with a bias towards the *cmr-*ON state shifted to a largely *cmr-*OFF state during murine infection, suggesting that the *cmr-*OFF state is under positive selection during infection. Collectively, our data reveal that Clade 5 strains have distinct growth properties, which may allow them to inhabit diverse ecological niches.

**Author Summary:** The Clade 5 strains of the *Clostridioides difficile* species are so phylogenetically divergent that they almost meet the threshold of being a distinct species. Although these strains are ubiquitously isolated from agricultural and environmental settings and an important source of zoonotic and community-acquired infections, it is unclear whether they have distinct phenotypic properties that allow them to colonize diverse hosts or persist in the environment. By combining a novel anaerobic time-lapse microscopy method with automated image analysis, we discovered that Clade 5 strains grow faster than strains from other *C. difficile* clades and that they frequently form long chains. These chaining properties are driven by the environmentally responsive expression of a non-canonical signal transduction system, which our analyses suggest is selected against during murine infection. Collectively, our analyses reveal that Clade 5 strains have distinct growth properties that may promote their persistence in the environment.

## Introduction

*Clostridioides difficile* is a leading cause of nosocomial infections in the United States, with approximately 500,000 new infections and 14,000 deaths being attributed to this organism annually [1, 2]. As an obligate anaerobe, *C. difficile* relies on its hardy, metabolically dormant spore form to survive outside the host and transmit disease to new hosts [3]. When *C. difficile* spores are ingested by susceptible hosts [4, 5], they germinate and outgrow into vegetative cells that subsequently colonize the colon. The vegetative cells secrete toxins that damage gut epithelial tissue [6], which triggers an inflammatory response that can cause disease pathologies ranging from mild diarrhea to pseudomembranous colitis and even death [2, 4]. *C. difficile* also causes recurrent infections in ∼20% of infections, which can lead to more severe disease symptoms [2, 7, 8].

*C. difficile*’s success as a pathogen may be related to its tremendous genetic diversity [9–11], with genomic analyses indicating that *C. difficile*’s core genome represents only ∼10-20% of its pan-genome [9, 10]. The plasticity of its “open” pan-genome likely helps *C. difficile* colonize the gastrointestinal tract of diverse animals, from mammals to invertebrates, and persist in environmental reservoirs like sewage and compost [12]. Indeed, the *C. difficile* species is so genetically diverse that it has been divided into five distinct phylogenetic clades based on multi-locus sequence typing (MLST) analyses, and the clades have been further subdivided into different ribotypes (RTs) or sequence types (STs) [10, 13].

Between these five clades, there are notable differences in geographic and host distributions. Clade 1 is the largest, most heterogeneous clade with the broadest geographic distribution [14, 15]. It includes over 200 STs, which can contain both toxin-producing and non-toxigenic strains, including the well-characterized, genetically tractable toxigenic strain 630 [9]. Clade 2 harbors epidemic-associated strains found within the ribotype 027 (RT027) lineage. Strains from this ribotype have been associated with outbreaks in hospitals, particularly in North America, due to their frequent resistance to fluoroquinolones [16–18]. These epidemic strains can also cause severe disease symptoms in part due to their production of three toxins: TcdA, TcdB, and CDT (binary toxin) [6]. Although RT027 strains have frequently been associated with “hypervirulence,” there is considerable phenotypic diversity within this lineage with respect to virulence, toxin production levels, flagellar motility, and sporulation [19–22]. Clade 3 strains are relatively uncommon and harbor a unique cell surface due to their lack of CwpV production [23], but phenotypic analyses of biofilm formation and motility suggest that they share similarities with Clade 2 strains like R20291 [24]. Clade 4 contains RT017 (ST37) strains that only encode a single toxin, TcdB (i.e., TcdA^−^CDT^−^), and are often clindamycin- and fluoroquinolone-resistant. RT017 strains have been associated with outbreaks in Europe and North America and are the most common strains found in Asia [25]. Clade 5 is the most genetically distant from the other 4 main *C. difficile* clades and is thought to have emerged before Clades 1-4 [26]. While the average nucleotide identity (ANI) for Clades 1-4 ranges between 97.1 - 99.8%, Clade 5 strains exhibit ANI values around 96%, which is close to the ANI value demarcation used by NCBI to define organisms of the same species [10, 27]. Thus, Clade 5 strains appear to be actively diverging from Clades 1-4 [10].

Clade 5 strains are an increasing problem in healthcare and agricultural settings because they can cause severe disease in humans and are commonly found in livestock, particularly pigs [12, 28]. While other *C. difficile* strains have been known to infect both humans and animals, only Clade 5 strains have been associated with zoonotic transmission from both animal-to-human and human-to-animal [28, 29]. The mechanisms underlying this bidirectional zoonotic transmission are poorly understood, but the increased carriage of antimicrobial resistance genes by Clade 5 strains may contribute to their ability to persist in agricultural and community settings [28, 30]. Thus, Clade 5 strains are of particular relevance from a One Health perspective [12, 31], especially since they frequently cause community-acquired infections [30] and are often detected in retail foods [32]. These observations highlight the importance of understanding the unique properties of this group of strains. Indeed, a recent genomic analysis suggests that RT078/ST11 strains within Clade 5 frequently carry zinc acquisition and homeostasis genes [11].

Despite numerous genomic analyses revealing the remarkable genetic diversity of *C. difficile* strains, relatively few studies have investigated the phenotypic diversity between strains from different clades. Clade-specific differences in colony morphology between Clade 5 strains relative to Clade 1-4 strains have been described in a limited set of analyses [26, 33], suggesting that differences in growth and/or cellular morphology may exist within clades. While differences in bulk growth rates between *C. difficile* strains have been reported [34], most phenotypic analyses have been conducted on a limited subset of strains within a given clade. Furthermore, systematic comparisons of the growth properties of different clades have only recently been described [35], while comparisons of their cell morphology have not been performed to date.

Here, we compare the growth properties of multiple strains derived from all five phylogenetic clades of *C. difficile* using anaerobic time-lapse microscopy. These analyses unexpectedly reveal striking differences in the growth and cell morphology of the Clade 5 lineage relative to strains from Clades 1-4. Specifically, we found that Clade 5 strains grow faster and frequently form long chains, in contrast with strains from Clade 1-4 strains. Genomic comparisons and genetic analyses indicate that the chaining phenotype of Clade 5 strains is driven by the phase-variable expression of the *cmrRST* operon by the invertible *cmr* switch [33]. Interestingly, we found that Clade 5 strains with a strong *cmr*-ON bias mostly reverted to a *cmr*-OFF phenotype during murine infection. Taken together, our data reveal that Clade 5 strains have unique growth properties relative to Clade 1-4 strains that may contribute to the widespread distribution of Clade 5 strain among diverse animal hosts.

## Results

### Development of a simple method for time-lapse imaging under anaerobic conditions

Time-lapse imaging of single cells has been widely used to study phenotypic heterogeneity in bacteria, which can impact important traits like antibiotic resistance and virulence [33, 36–39]. However, live single-cell analyses in *C. difficile* have been complicated by its inability to grow in the presence of atmospheric oxygen [40]. While time-lapse microscopy analyses of *C. difficile* have previously been reported, they require custom growth chambers to maintain anaerobic conditions [41] and thus may limit the accessibility of these experimental systems to a broad range of investigators.

To overcome these limitations, we established a simple system that relies solely upon commercially available reagents and materials to grow *C. difficile* cells under anaerobic conditions. This system uses gas-tight, adhesive Gene Frames, which have been used extensively in imaging applications for bacteria [42]. Notably, the gas-impermeability of these commercial seals allows anaerobic conditions to be maintained when agarose pads made with growth media are prepared in the anaerobic chamber (**Figure 1**). Gene Frames also generate thick agarose pads, which are critical for *C. difficile* to grow in a sealed system under ambient conditions. After agarose pads are prepared in the anaerobic chamber, *C. difficile* cultures are inoculated onto the pads, and the pads are sealed with a coverslip. The growth chamber is then removed from the chamber and imaged on a heated microscope stage under ambient conditions for up to 6 hours or until *C. difficile* stops growing as a monolayer.

**Figure 1.**
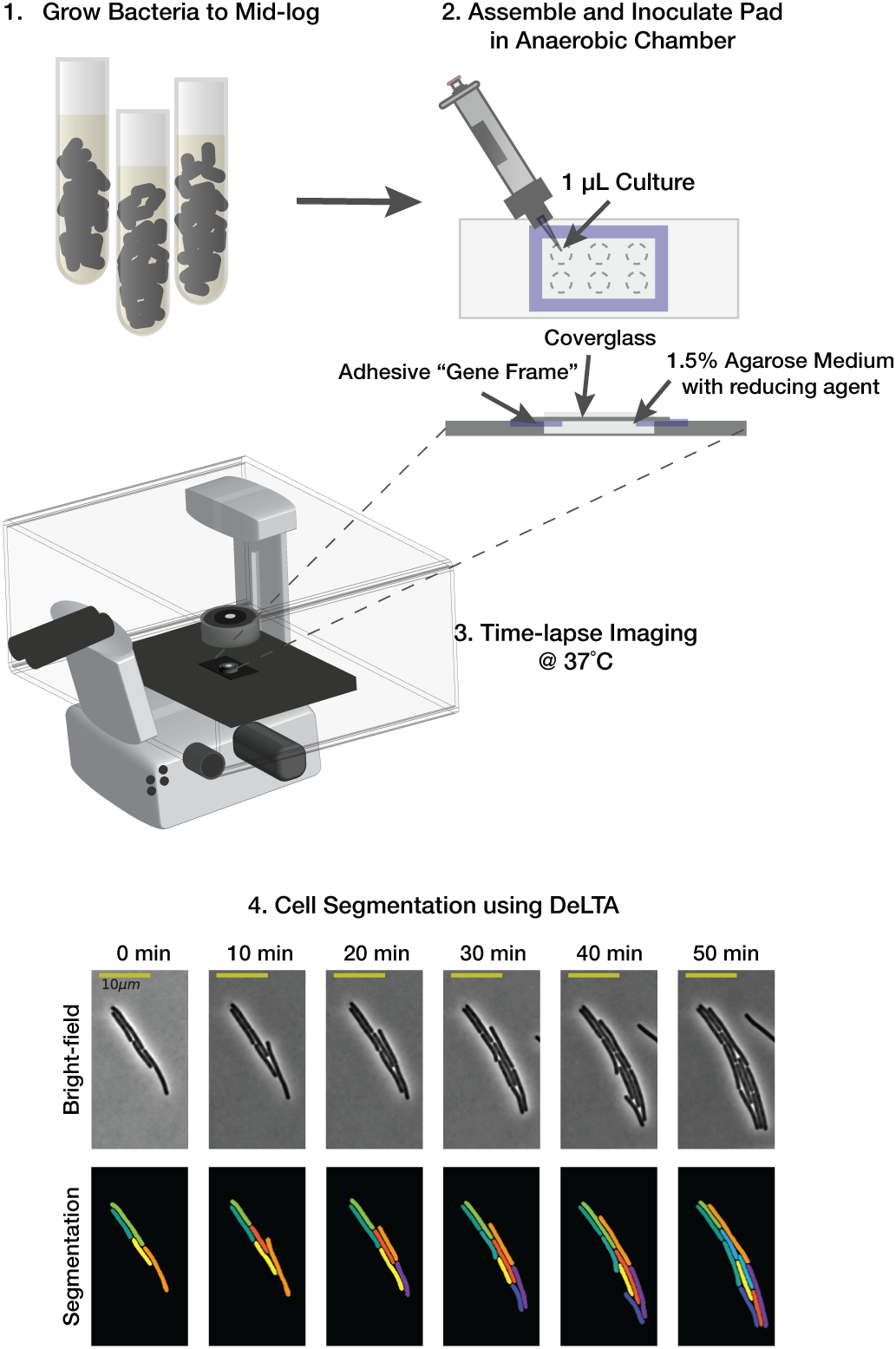
Schematic of the anaerobic single-cell imaging set-up. Exponentially growing *C. difficile* cells in TY medium supplemented with cysteine (TYC) are spotted onto 1.5% agarose pads formed within gas-tight adhesive Gene Frames inside the anaerobic chamber. Up to 6 strains can be spotted onto a pad. The pad is sealed with a coverslip, and the imaging chamber is removed from the anaerobic chamber and transferred to a heated (37°C) microscope stage. Time-lapse microscopy is used to visualize the growth of individual bacterial cells for 2-6 hrs. The output data is segmented and tracked with the DeLTA Python package (191). An example filmstrip of strain 630 grown on TYC medium over time is shown (Bottom).

### Time-lapse microscopy reveals clade-specific differences in elongation rate and cell length

Having established an anaerobic time-lapse imaging setup, we compared the single-cell growth properties of representative *C. difficile* strains from Clade 1 (630, ribotype (RT) 012), Clade 2 (R20291, RT027), Clade 3 (E15, RT075), Clade 4 (M68, RT017), and Clade 5 (M120, RT078) (**Figure 2**, **Table 1**). The five “representative” strains were all isolated from patients with *C. difficile*-associated disease and are frequently used as reference genomes for their clades and ribotype groups. Notably, RT027 (ST1), RT017 (ST45), and RT078 (ST11) strains are from ribotypes/ multi-locus sequencing types that are frequently isolated from patients with *C. difficile* infection (CDI) [12, 16, 19, 25]. In contrast, Clade 3 strains are more rare and the least characterized of *C. difficile* strains [23].

**Figure 2.**
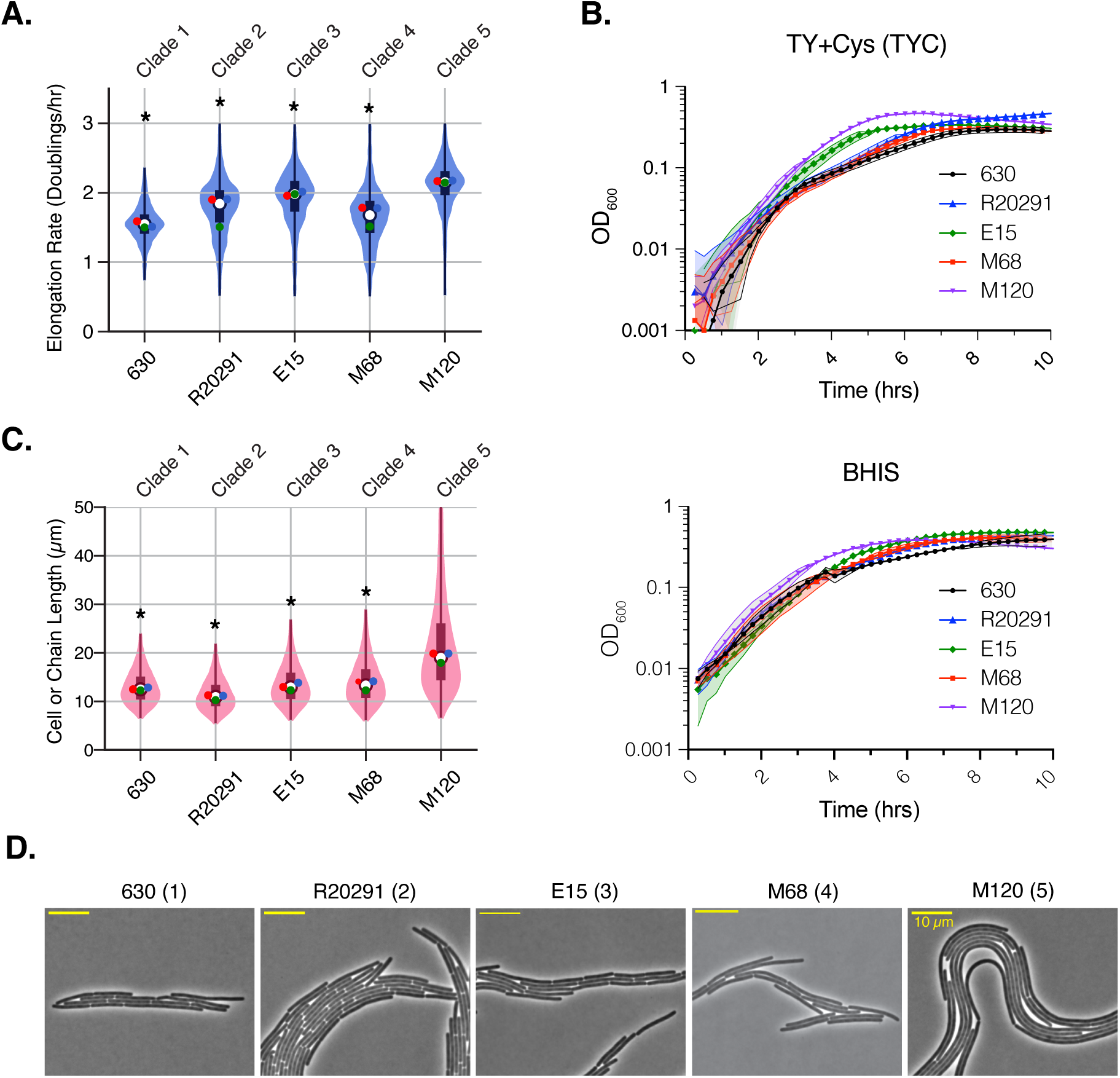
Clade 5 strain M120 elongates more quickly and exhibits cell chaining. (A) Violin plot of the elongation rates measured during time-lapse microscopy analyses of strains 630 (Clade 1), R20291 (Clade 2), E15 (Clade 3), M68 (Clade 4), and M120 (Clade 5) grown on TY supplemented with cysteine (TYC) agar. Data are from three biological replicates, with the mean of each replicate shown as a point on the violin. (B) Optical density-based analyses of bulk population growth of the indicated strains in TYC or BHIS media. The number in brackets indicates the clade to which a given strain belongs. (C) Violin plot of the cell or chain length measured during time-lapse microscopy for the strains shown in **A**. Each replicate mean is shown as a point on the violin. Statistical significance for A and B was determined by comparing the mean of the three replicates of strains from Clades 1-4 strains relative to the Clade 5 M120 strain using a Kruskal-Wallis test * p < 0.05. (B) Phase-contrast image from time-lapse microscopy movies. Scale bar is 10 μm.

**Table 1.** *Clostridioides difficile* clinical isolates used in this study.

| Strain Name | Clade | Ribotype | ST group | Source | Reference |
| --- | --- | --- | --- | --- | --- |
| 630 | 1 | 012 | 2 | Zurich, 1982 (Sanger Institute) | [87] |
| BBL2 | 1 | 012 | 2 | Memorial Sloan Kettering | [20] |
| WU38 | 1 | 012 | 2 | Barnes-Jewish Hospital | [20] |
| 190B | 1 | 087 | 46 | Memorial Sloan Kettering | [20] |
| R20291 | 2 | 027 | 1 | London, 2006 (Sanger Institute) | [88] |
| Wup14 | 2 | 027 | 1 | Barnes-Jewish Hospital | [20] |
| BBL4 | 2 | 027 | 1 | Memorial Sloan Kettering | [20] |
| 186A | 2 | 027 | 1 | Memorial Sloan Kettering | [20] |
| E15 | 3 | 075 | unknown | France (Bruno Dupuy/ Lynn Bry) | [89, 90] |
| BI1 | 3 | unknown | unknown | Bruno Dupuy/ Lynn Bry | This study |
| 95-978 | 3 | unknown | unknown | Bruno Dupuy/ Lynn Bry | This study |
| M68 | 4 | 017 | 81 | Dublin, 2006 (Sanger Institute) | [53] |
| 1002 | 4 | unknown | 39 | Memorial Sloan Kettering | [20] |
| M120 | 5 | 078 | 11 | UK, 2007 (Sanger Institute) | [53] |
| TAL28131 | 5 | 078 | 11 | NY Presbyterian/Weill Cornell Medical Center | [91] |
| TAL29600 | 5 | 078 | 11 | RM Alden Research Lab | [91] |
| TAL29996 | 5 | 078 | 11 | Vines VA Hospital | [91] |
| TAL30550 | 5 | 078 | 11 | Mayo Clinic | [91] |
| TAL30574 | 5 | 078 | 11 | Tufts Medical Center | [91] |
| V48 | 5 | 078 | 11 | Brigham & Women's Hospital | This study |
| 139b | 5 | 078 | 11 | Memorial Sloan Kettering | [20] |
| WU66 | 5 | 078 | 11 | Barnes-Jewish Hospital | [20] |

Memorial Sloan Kettering Cancer Center (MSK), Barnes-Jewish Hospital (BJH)

The growth properties of single cells visualized by time-lapse microscopy were quantified using Deep Learning for Time-lapse Analysis (DeLTA) software, which rapidly and accurately segments and tracks bacteria growing in two dimensions on agarose pads [43, 44]. This software uses deep convolutional neural networks to analyze time-lapse microscopy images, allowing the growth properties of individual cells growing in microcolonies on agarose pads to be determined. The segmentation and tracking of *C. difficile* cells were highly accurate (**Figure 1**), and minimal user input or post-image processing was needed to obtain growth property measurements.

Robust growth was observed for all strains using our system. Growth was quantified by measuring the elongation rate, which was defined as doublings/hr to indicate the number of times that a cell’s length doubles in one hour. The elongation rate (doublings/hr) is distinct from the doubling time, or generation time, which represents the length of *time* that it takes before a bacterium divides. Instead, the elongation rate reflects how fast the cell is increasing in length over time. Notably, the Clade 5 strain M120 elongated the fastest (2.1 doublings/hr, p < 0.05), followed by Clade 3 E15 (2.0 doublings/hr) and Clade 2 strain R20291 (1.8 doublings/hr), and then Clade 1 strain 630 and Clade 4 strain M68 (1.6-1.7 doublings/hr) (**Figure 2A**).

Importantly, the differences in single-cell elongation rates measured for the five strains were also observed in bulk population analyses of their growth using optical density in TYC and BHIS media (**Figure 2B** & **S1**). These analyses confirmed that the Clade 5 strain M120 grew faster (based on optical density-based analyses) in these media than the Clade 1-4 strains (p < 0.001). In contrast, negligible differences in bulk growth rates were observed between Clade 1-4 strains in BHIS media, although, in TYC medium, the Clade 3 strain grew faster than Clades 1, 2, and 4 strains (**Figures 2B** & **S1**).

The Clade 5 strain M120 exhibited another distinct growth property from the Clade 1-4 strains. While strains from Clades 1-4 produced cells of similar length prior to cell division, with an average apparent length of ∼13 µm, cells of Clade 5 strain M120 were significantly longer, with an average apparent length of ∼22 µm. Indeed, cells ∼50 µm were readily observed for the Clade 5 strain M120 (**Figure 2C**), and these cells appeared to bend readily (**Figure 2D**). By incorporating the FM4-64 membrane stain into the agarose pads to visualize division septa [45], we assessed whether the Clade 5 strain M120 forms chains vs. filaments, These analyses revealed that septa were readily observed in Clade 5 strain M120 across the length of a given cell (**Figure 3, inset**), Since the spacing between division septa was relatively consistent, the Clade 5 strain M120 appear to undergo cell separation less efficiently than strains from the other clades tested. Indeed, cell separation in strain M120 was so inefficient that it was often necessary to stitch together several fields of view to fully visualize M120 chains, which approached several hundred microns and even up to ∼1 mm in length (**Figure 3**).

**Figure 3.**
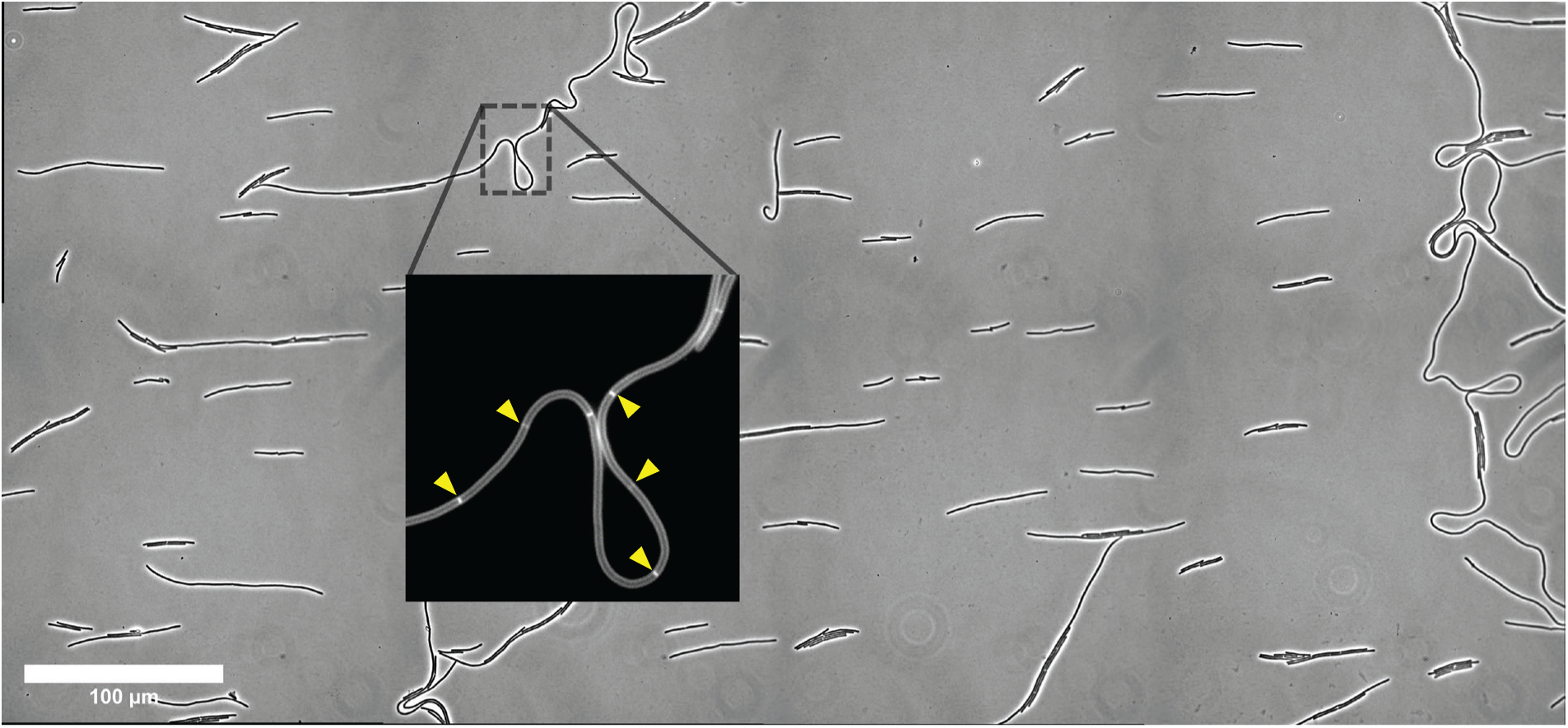
*C. difficile* Clade 5 can form large heterogenous chains. Large mosaic phase-contrast image of the Clade 5 strain M120. Inset shows chains revealed by staining with the membrane dye FM4-64; septa are highlighted with yellow arrows. The image was stitched from 8 individual fields of view at 63X magnification. Scale bar is 100 μm.

### Clade 5 clinical isolates typically form long chains and grow more quickly than strains from other clades

Since prior work indicated that Clade 5 strains produce colony morphologies distinct from Clade 1, 2, and 4 strains [26], we sought to determine whether the striking cell chaining phenotype and faster growth rate observed in Clade 5 strain M120 were properties shared by other Clade 5 strains. Thus, we compared the single-cell growth properties of five additional Clade 5 clinical isolates obtained from several hospitals around the country on TYC agarose (**Table 1**) using time-lapse microscopy analyses (**Figure 2**). Clade 1 strain 630 was included as a control since it does not form chains in any of the conditions we have tested.

These analyses revealed that all but one of the Clade 5 strains tested formed long chains, with TAL29600 forming the longest chains (29 µm on average, **Figure 4A-C**). In contrast, strain TAL29996 formed shorter chains that were comparable in length to those observed for strains from Clades 1-4 (12-13 µm, **Figures 2B** & **4C**). Notably, DeLTA segmented many Clade 5 chains as single cells because cell separation (i.e., invagination) had not yet initiated at division septa visualized via FM4-64 staining. To overcome this limitation and accurately quantify cell length within long chains, namely the distance between division septa, we modified our image processing pipeline to use a thresholding method to detect division septa. After generating masks in DeLTA to segment the chains, we modified the mask so that only the interior of the contour was analyzed. We then applied an adaptive thresholding method to identify division septa based on their elevated fluorescence relative to the long axis of the cell; the thresholds were defined using a Gaussian-weighted method (**Figure 4A**, red dots, **Figure 4B**). Septa detected with this automated method were also manually inspected. These analyses revealed that the average length of cells within the chains of cells made by Clade 5 strains is only slightly longer than the average length of cells made by strains that do not form chains, namely Clade 1 strain 630 and Clade 5 strain TAL29996 (**Figure 4C**, **Table 2**). For example, the Clade 5 strains that produced ”he l’ngest chains (TAL29600 and TAL30550) had average cell lengths of 14 and 12 µm, respectively, which is only 30-50% longer than the non-chaining strains 630 (Clade 1, 8.5 µm) and TAL29996 (Clade 5, 9.4 µm) (**Table 2**). Notably, even though the Clade 5 strain TAL29996 did not form chains, it still exhibited higher elongation rates, which were similar to those measured for other Clade 5 strains (2.1 doublings/hr vs. 1.7 doublings/hr for Clade 1 strain 630, **Figure 4D**).

**Figure 4.**
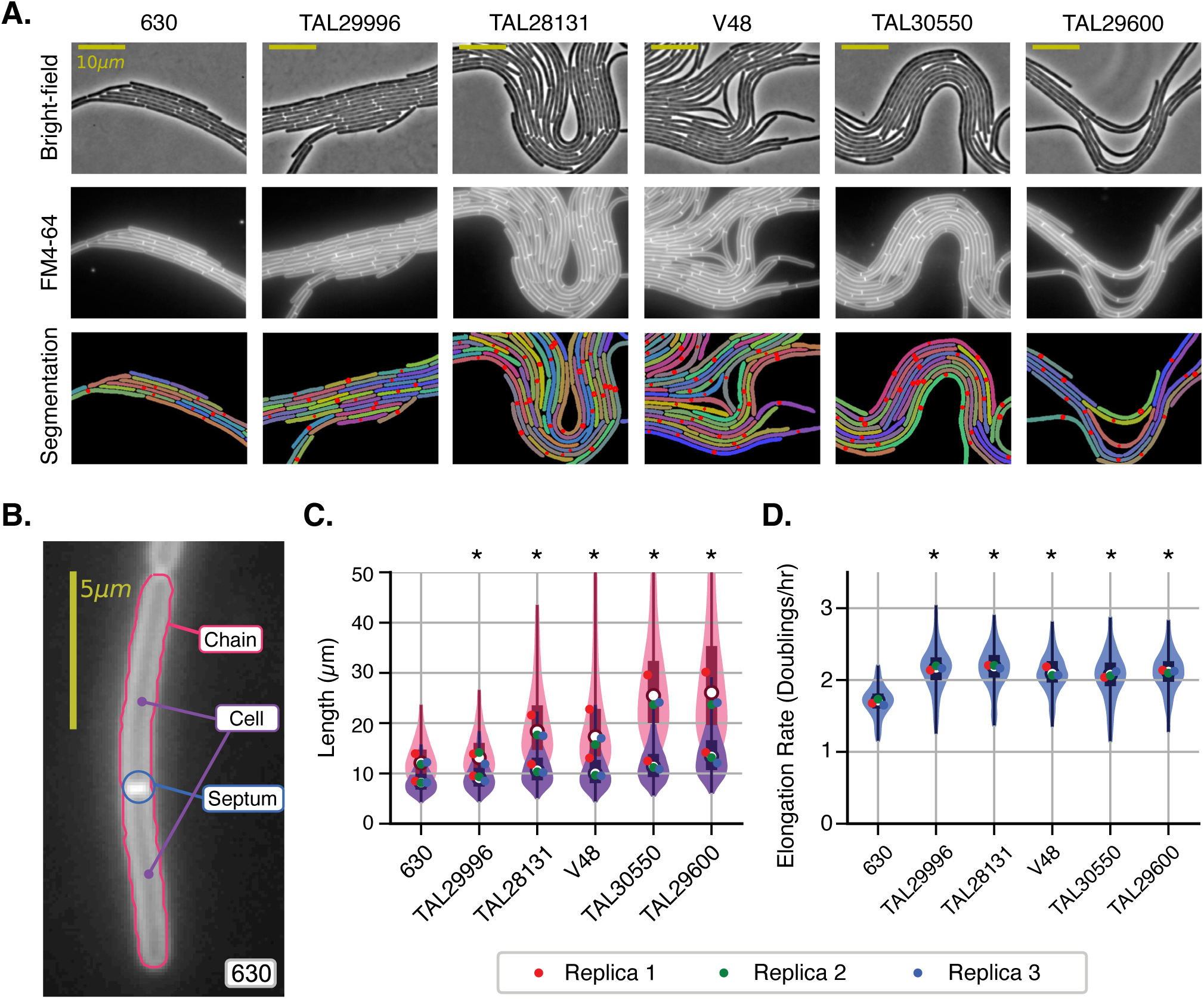
Faster growth is a common feature of Clade 5 strains, but the chaining phenotype is not fully penetrant in Clade 5 strains. (A) Phase-contrast microscopy images from time-lapse microscopy studies of Clade 5 strains; Clade 1 strain 630 is included for comparison. Fluorescence microscopy was used to visualize the FM4-64 stain incorporated into the agarose pads. Masks generated using DeLTA are shown, with the bottom panel showing division septa identified with our adaptive thresholding approach (red dots). (B) Example image showing the parameters identified using DeLTA combined with our adaptive threshold method for detecting division septa (red dot) within a chain of cells. Following the automated thresholding analysis for detecting septa, the images were manually inspected to ensure that all septa were properly identified. (C) Cell or chain length measured using automated DeLTA analyses (pink violin plot) or DeLTA combined with the adaptive threshold-manual inspection analyses (purple violin plot). The former method is more likely to measure chain length, while the latter method accurately measures cell length. (D) Violin plot of the elongation rates measured based on three biological replicates. Each replicate mean is shown as a point on the violin; statistical significance was determined by comparing the mean of the three replicates using a Kruskal-Wallis test, * p < 0.05.

**Table 2:** Cell size and growth rate statistics for the studied strains.

| Strain Name | Clade | Mean Growth rate | Mean Cell Length (um) | Mean Chain Length (um) |
| --- | --- | --- | --- | --- |
| 630 | 1 | 1.63 ± 0.02 | 8.5 ± 2.2 | 12.5 ± 3.5 |
| R20291 | 2 | 1.88 ± 0.01 | not measured | 11.7 ± 3.2 |
| E15 | 3 | 1.92 ± 0.01 | not measured | 13.6 ± 4.2 |
| M68 | 4 | 1.64 ± 0.01 | not measured | 13.7 ± 4.3 |
| M120 | 5 | 2.15 ± 0.01 | not measured | 21.8 ± 10.2 |
| TAL29996 | 5 | 2.15 ± 0.02 | 9.4 ± 2.7 | 13.7 ± 3.9 |
| TAL28131 | 5 | 2.17 ± 0.02 | 11.2 ± 3.3 | 19.7 ± 6.9 |
| V48 | 5 | 2.10 ± 0.02 | 10.8 ± 3.7 | 19.4 ± 8.5 |
| TAL30550 | 5 | 2.05 ± 0.04 | 11.8 ± 3.3 | 27.3 ± 9.8 |
| TAL29600 | 5 | $2.11 \pm 0.02$ | $14.0 \pm 4.4$ | $29.1 \pm 13.1$ |

The ± interval represents the 95% confidence interval of the mean growth rate. The standard deviation (S.D) is provided.

Importantly, the additional Clade 5 strains tested also grew faster in bulk optical density-based analyses in broth culture than Clade 1-4 strains irrespective of their ability to form chains (**Figure S2**). To assess whether these findings would extend to additional Clade 5 strains vs. Clade 1-4 strains, we analyzed the growth of additional strains from all five clades in broth culture. These analyses confirmed that Clade 1-4 strains grow at similar rates in BHIS media, which are slower than those observed for the nine Clade 5 strains analyzed in this media (**Figure S2**). However, Clade 3 strains grew relatively faster than the Clade 1 630 strain in TYC medium, but their growth was still slower than the Clade 5 strain M120 (**Figure S2**). Taken together, these analyses strongly suggest that Clade 5 strains grow faster than Clade 1-4 strains and are more likely to form chains, presumably because their cell separation mechanisms are less efficient.

### Cell chaining in Clade 5 strains is not dependent on growth on a solid medium

*C. difficile* has previously been shown to promote cell elongation and chain formation upon induction of the *cmrRST* locus [33], which encodes a non-canonical signal transduction system. Expression of this locus is also responsive to c-di-GMP levels [46], which increases in cells grown on solid surfaces such as in a biofilm or on an agar plate [46]. To test whether the chaining phenotype observed in Clade 5 strains is induced by growth on a surface, we assessed the chaining properties of Clade 5 strains during logarithmic growth in rich medium broth culture using the fluorescent D-amino acid label, HADA, to stain septa. These analyses revealed that Clade 5 strains still form chains during broth culture growth, although the chains are not as long as those observed during growth on the agarose pads (**Figures 5 and S3**).

**Figure 5.**
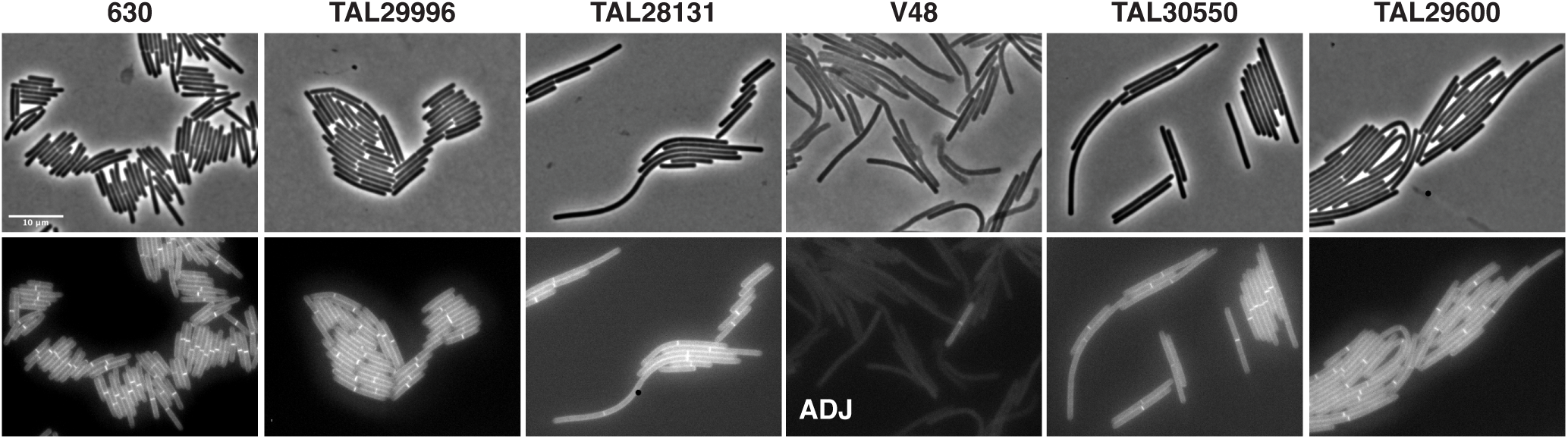
Clade 5 strains form chains during logarithmic growth in broth culture. Representative micrographs showing phase-contrast (top) and peptidoglycan labeling with the fluorescent D-amino acid, HADA, (bottom) following growth in rich broth (BHIS) to mid-logarithmic phase. All strains shown, with the exception of Clade 1 strain 630, are Clade 5 strains. Scale bar, 10 µm. Data are representative of three independent experiments. **ADJ** indicates that the brightness of the image was enhanced to detect HADA labeling in V48.

Since Clade 5 strains grow rapidly in rich media, we considered the possibility that the chaining phenotype might be mitigated by allowing more time for cell separation to occur after cell division. To test this possibility, we grew Clade 5 strains in CDDM minimal medium and analyzed their chaining properties [47]. While Clade 5 strains grew slower in CDDM medium relative to richer media (compare **Figure S4A** to **Figures 2 & S2**), the Clade 5 strains nevertheless formed chains in minimal medium (**Figure S4B**), with the exception of strain TAL29996 strain. Taken together, our results reveal that Clade 5 strains undergo cell separation less efficiently in a range of growth conditions relative to strains from other clades.

### Cell length does not correlate with the propensity to sporulate

We next wondered whether the propensity to form chains impacts the ability of Clade 5 strains to sporulate. Analyses in *Bacillus subtilis* suggest that smaller cells, such as those formed during stationary phase growth [48], are more likely to sporulate likely because they concentrate proteins involved in the phosphorelay that induces sporulation [49, 50]. For example, a decrease in cell length in *B. subtilis* helps the kinase KinA reach the threshold concentration needed to trigger sporulation initiation. Although *C. difficile* lacks homologs of KinA and other components of the phosphorelay system [51], the longer cells generated by Clade 5 strains may be less likely to induce sporulation due to dilution of a currently unknown sporulation regulator. To test this hypothesis, we analyzed the propensity of Clade 5 strains to form spores when plated on 70:30 sporulation medium using phase-contrast microscopy and heat resistance assays. These analyses revealed that Clade 5 strains exhibit striking differences in sporulation frequency, with some strains exhibiting close to 100% sporulation levels, and others exhibiting levels closer to 20% (**Figures 6 & S5**). Interestingly, strain TAL29600 exhibited extremely low levels of sporulation (0.003%), and it continued to form chains during growth on 70:30 medium. In contrast, the other Clade 5 strains analyzed did not form long chains when grown on 70:30 sporulation medium, and the spores produced by these isolates exhibited similar lengths and proportions relative to the Clade 1 strain 630 (**Figure S6**). These data suggest that Clade 5 strains alter their cell length and propensity to form chains depending on the growth conditions encountered.

**Figure 6.**
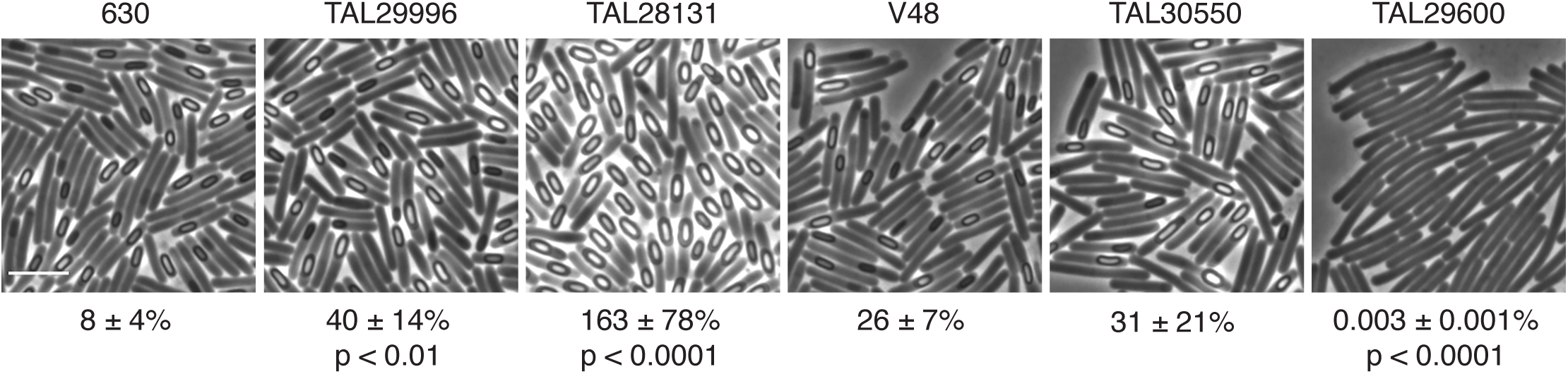
Sporulation levels in Clade 5 isolates grown on 70:30 medium. Phase-contrast microscopy of the indicated strains ∼24 hrs after sporulation induction. All strains shown with the exception of 630 (Clade 1) are Clade 5 strains. The percent heat-resistant spores is indicated below the respective images. The percentage was determined from 20-24 hr sporulating cultures and represent the mean and standard deviation for a given strain based on a minimum of three biological replicates. Statistical significance relative to strain 630 was determined using a one-way ANOVA and Tukey’s test. The scale bar represents 5 µm.

Consistent with this hypothesis, even though strain M120 forms long chains during growth in or on rich medium (**Figures 3** & **S3**), it readily formed spores during growth on 70:30 medium (∼100% sporulation frequency, **Figure S5**). Notably, the average cell length for visibly sporulating Clade 5 cells was ∼5-6 µm (**Figure S8**) compared to the average ∼11 µm cell length measured for the Clade 5 strains in rich broth culture. This reduced cell length was observed even for Clade 5 strain TAL29600 (**Figure S8**), which sporulates poorly on 70:30 medium, indicating that there was little correlation between cell length and propensity to sporulate for the Clade 5 strains analyzed.

### Comparative genomics reveals that the chaining phenotype of Clade 5 strains is driven by *cmrRST* operon expression

Given the phenotypic difference in chaining observed for the TAL29996 strain relative to the 8 other Clade 5 strains analyzed, we sought to gain insight into the mechanism driving this difference by comparing the genomes of five of the Clade 5 strains, including TAL29996. These analyses revealed that the average nucleotide identity (ANI) for orthologous genes ranged between 99.83-99.99% (**Table S1**) and that the pan-genome between the five strains is 12%. Thus, all five strains are quite closely related. The pan-genome analysis revealed that, relative to the other four strains, TAL29996 is missing one duplication of *blaR1*, which encodes an integral membrane protein that senses beta-lactams, and a gene region predicted to be involved in nicotinate metabolism. To identify SNPs that might distinguish TAL29996 from the other strains, we used breseq [52]; the Clade 5 strain M120 genome sequence was used as the reference genome because it is the Clade 5 strain traditionally characterized [34, 53]. These analyses identified 10 SNPs that were unique to TAL29996, but none were obviously involved in regulating cell separation or peptidoglycan synthesis (**Table S2**).

We next took a candidate approach to gain insight into why TAL29996 mediates cell separation more efficiently than the other Clade 5 strains by analyzing the orientation of the *cmr* switch, also known as the Cdi6 DNA invertible element [33, 54]. This phase-variable element affects the expression of the adjacent *cmrRST* operon, which encodes a non-canonical CmrRST signal transduction system that regulates cell chaining and colony morphology [33, 46, 54]. Cells from rough colonies form chains and are highly biased to the ON orientation of the *cmr* switch, whereas cells from smooth colonies do not form chains and are highly biased to the OFF orientation [33]. Although the sequence of the *cmr* switch is identical between the Clade 5 strains analyzed, including strain TAL29996, we found that the orientation of the *cmr* switch during growth in broth culture was markedly different for Clade 5 strain TAL29996. Specifically, qPCR analyses revealed that Clade 5 strains that form chains are biased towards the *cmr*-ON orientation (between 70-96% ON) (**Figure 7A**), whereas the *cmr-*ON orientation was markedly less frequent in the non-chaining strain TAL29996 (∼20%, **Figure 7**).

**Figure 7.**
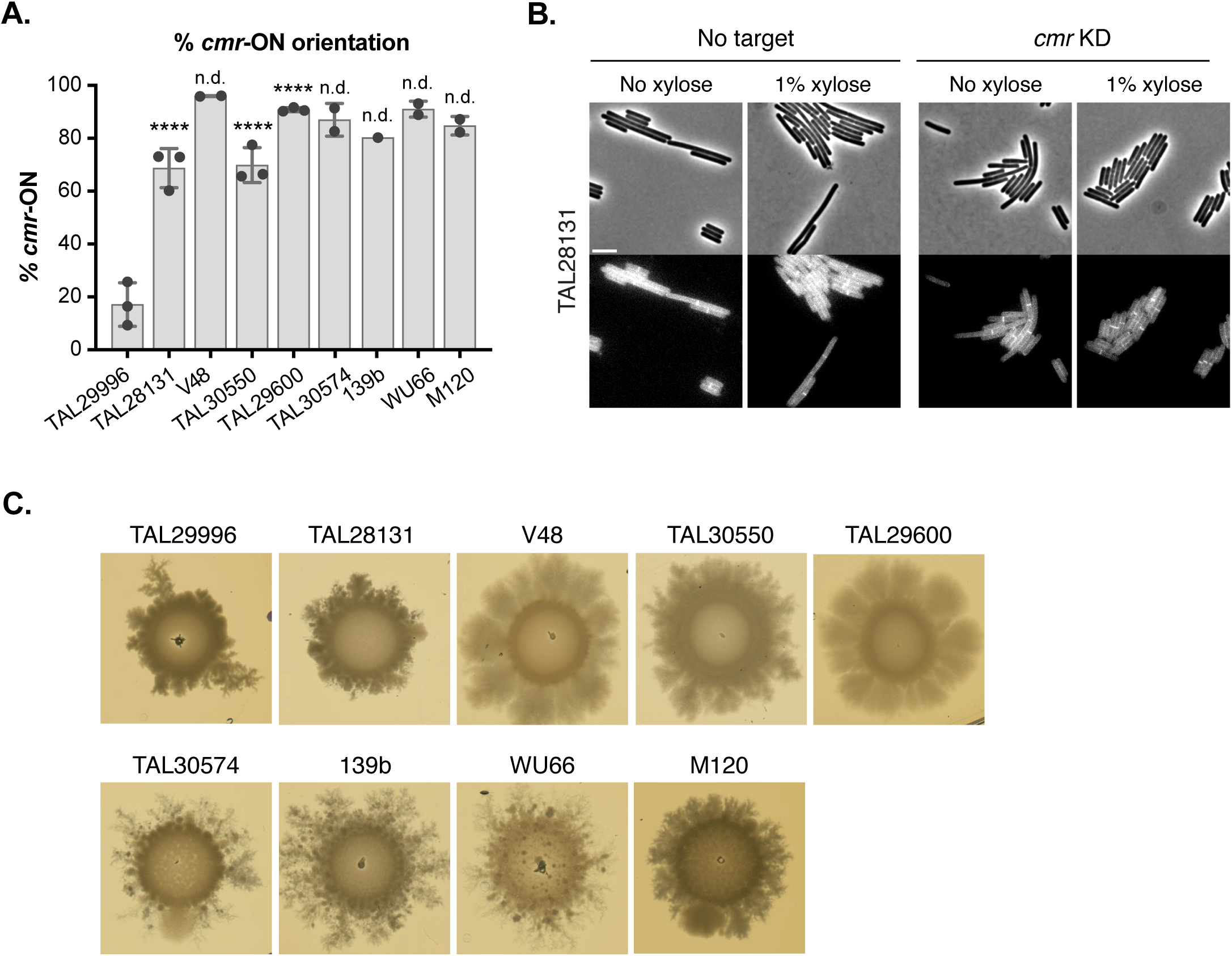
The orientation of the *cmr* switch promotes Clade 5 strain cell chaining and surface motility. (A) Orientation-specific qPCR for detecting the orientation of the *cmr* switch in the indicated strains. The mean and standard deviation based on one to three biological replicates are shown. Statistical significance relative to strain TAL29996 was determined using a one-way ANOVA and Tukey’s test for strains where data from three independent experiments was obtained. **** p < 0.0001. (B) Representative micrographs of *cmrRST* CRISPRi knock-down strains compared to a no target control. Phase-contrast (top) and peptidoglycan labeling with the fluorescent D-amino acid, HADA, (bottom) images following growth in rich broth (BHIS) to mid-logarithmic phase. Scale bar, 5 µm. (C) Representative images of surface motility 5 days after exponentially growing liquid cultures of the indicated strains were spotted onto BHIS agar plates.

Since these analyses correlated the *cmr*-ON switch orientation to the chaining phenotype of Clade 5 strains, we tested whether knocking down the expression of the *cmrRST* operon using CRISPRi in strains that are biased towards the *cmr*-ON state would reduce their chaining properties. A plasmid targeting the *cmrR* gene using CRISPRi was introduced into the Clade 5 strains TAL28131, TAL30550, and TAL30574, which typically produce chains and are found predominantly in the *cmr-*ON state in rich broth culture (70-96%). Knocking down the expression of the *cmrRST* operon in all three strain backgrounds reduced their chaining phenotypes under these conditions (**Figures 7B** & **S8**), indicating that the expression of the *cmrRST* operon in Clade 5 strains drives their propensity to form chains.

### Surface motility in Clade 5 strains relative to strains of other clades

Since the *cmr-*ON state has also been correlated with increased surface motility [33], we analyzed the surface motility of our Clade 5 strains. Consistent with prior reports [33], the primarily *cmr-*ON state strains exhibited greater and more uniform surface motility (**Figure 7C**), whereas the predominantly *cmr-*OFF state TAL29996 strain exhibited less and more asymmetric surface motility, with fractal-like extensions emerging from only a few sites. This asymmetric phenotype has previously been reported for the Clade 2 strain R20291, whose *cmr* switch is predominantly in the OFF position in liquid cultures but converts to the ON orientation during growth on plates [33]. These observations suggest that, even though TAL29996 is biased to the *cmr*-OFF orientation during broth culture growth, a subset of TAL29996 cells switch to the *cmr*-ON orientation during growth on BHIS agar, leading to the asymmetric spreading phenotype.

We next assessed whether additional Clade 1-4 strains exhibit surface motility. These analyses revealed that asymmetric motility was more frequently observed in Clade 2 strains, although Clade 2 strain Wup14 exhibited little surface motility (**Figure S9**). Clade 1 strains exhibited a range of surface motility, from high surface motility with strain 630 to lower surface motility with strain WU38 (**Figure S9**). While the data suggest that *cmr* switching in Clade 2 strains varies between strains during growth on agar medium, Clade 5 strains biased towards the *cmr-*ON state are more likely to exhibit surface motility. However, it is important to note that additional factors contribute to surface motility beyond expression of the *cmrRST* operon [33], since loss of pili can also decrease surface motility on plates [55]. Regardless, the data imply that the *cmr*-ON state promotes surface motility in Clade 5 strains.

### Colonization and virulence properties of Clade 5 strains

Beyond the effects of the CmrRST system on cell chaining and surface motility, this system has also been shown to impact the virulence of the Clade 2 strain, R20291, in a hamster model of infection, with loss of *cmrR* reducing R20291’s ability to cause disease and the *cmr*-OFF orientation correlating with less severe disease in hamsters [33]. Since chaining in *Bacillus anthracis* strains promotes virulence [56], while chaining in *Enterococcus faecalis* promotes colonization [57], we compared the ability of Clade 5 strains to colonize mice and cause disease. Mice were infected with 10^5^ spores of several Clade 5 strains and the Clade 1 strain 630 and the weight loss induced by these strains, their colonization levels, and orientation of the *cmr* switch over the course of the 14-day infection were assessed. For this latter analysis, we focused on strains TAL29600 and TAL29996 because they exhibited the highest and lowest *cmr*ON orientations, respectively, during growth in rich media (**Figure 7A**).

All Clade 5 strains tested colonized to relatively similar levels throughout the 14 days of the infection. Strain 630 also colonized mice to similar levels in the first two days of infection and then maintained colonization, albeit at 1-2 logs lower than the Clade 5 strains (**Figure 8A**). Interestingly, only the Clade 5 strain TAL29600 caused significant weight loss relative to the other strains on Days 2 through 4, although the Clade 1 strain 630 caused some weight loss on Day 3 (**Figure 8B**). This latter phenotype is consistent with prior reports of strain 630 causing only mild disease symptoms in the cefoperazone model of murine infection [58, 59]. Analyses of the *cmr* orientation revealed that the *cmr-*OFF orientation appeared to be selected for over the course of the infection. While the TAL29600 spore inoculum started off with ∼30% *cmr-*ON frequency, the frequency of TAL29600 cells detected in the *cmr-*ON orientation decreased rapidly to < 5% *cmr*-ON by 24 hrs post-inoculation (**Figure 8C**). As the infection progressed, two of the 8 mice tested exhibited an increase in TAL29600 cells with the *cmr*-ON orientation (6-30% *cmr-*ON) (**Figure S10**). Conversely, the TAL29996 strain retained the ∼1% *cmr*-ON frequency of the inoculum for the greater part of the 14-day infection (**Figure 8C**). Taken together, these analyses reveal that Clade 5 RT078 strains efficiently colonize mice but vary in their ability to cause disease. Furthermore, the ability of the Clade 5 strains to colonize or cause disease did not strongly correlate with their ability to form chains in broth culture.

**Figure 8.**
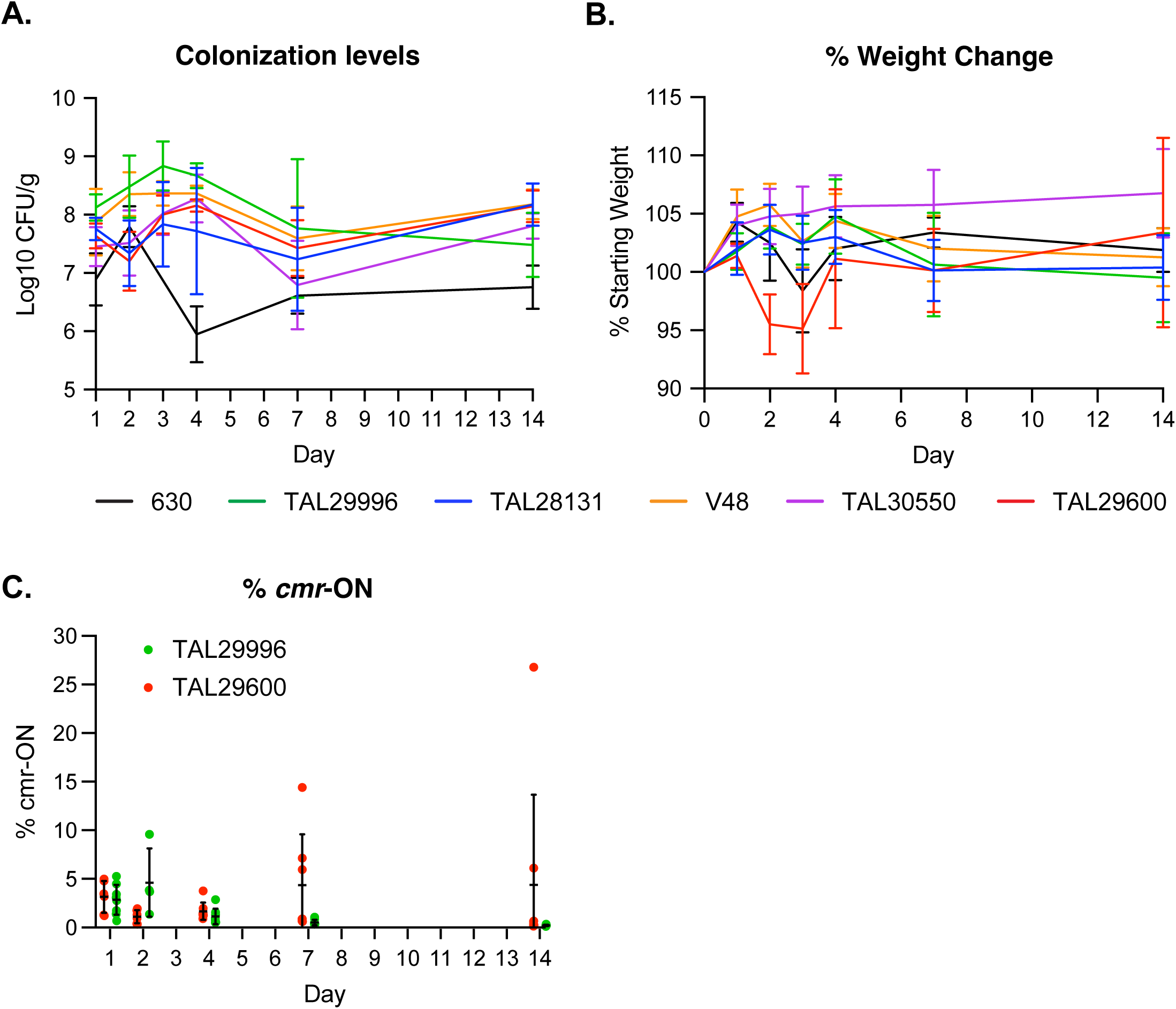
Infection and colonization dynamics of Clade 5 strains in mice. (A) Fecal colony-forming units measured by selective plating and (B) Percentage of weight loss to baseline of infected mice on Days 1-4, 7, and 14. The mean and standard deviation are shown based on the results of two experiments consisting of four mice each (n = 8). (C) Percentage of *cmr*-ON switch orientation measured in fecal pellets on the indicated days by qPCR. For the strain TAL29996 inoculum, 1% of the spores had the *cmr*-ON switch orientation, while the TAL29600 inoculum consisted of 32% *cmr*-ON spores. The mean and standard deviation based on analyses of eight mice are shown, although fecal pellets could not be collected from some of the mice on Day 2. The same mouse exhibited a higher *cmr*-ON frequency for TAL29600 over time (15%, Day 7 and 27%, Day 14).

## Discussion

While Clade 5 strains are genetically distinct [10, 26] and more prevalent in animals than Clade 1-4 strains [12, 29, 30], the phenotypes that distinguish Clade 5 strains from Clade 1-4 strains are not well understood. By phenotypically characterizing *C. difficile* strains from multiple clades using time-lapse microscopy, we discovered that Clade 5 strains have distinct growth properties from Clade 1-4 strains. Specifically, Clade 5 strains elongate more quickly (**Figures 2, 4, S1**) and form long chains more readily than strains from Clades 1-4, irrespective of media type or growth on a surface (**Figures 2, 4, S2**, and **S4**). In contrast, long chains were not observed in any of the Clade 1-4 strains, regardless of whether the cells were grown on agarose pads or in broth culture. Thus, Clade 5 strains undergo cell separation far less efficiently than strains from other clades during growth in rich media. While our analyses of Clade 5 strains were largely limited to the RT078 ribotype, we note that a prior report described an RT126 strain isolated from a patient experiencing multiple recurrences that also formed long chains [60]. Since the RT126 ribotype is closely related to the RT078 ribotype [12], it is likely that the chaining phenotype will be observed in other Clade 5 ribotypes.

Our data indicate that the chaining phenotype of Clade 5 strains relates to their propensity to express the *cmrRST* operon [33, 46] because they favor the *cmr-*ON state (**Figure 7**). The non-chaining TAL29996 strain has a *cmr*-ON orientation frequency of 20%, whereas the 8 other Clade 5 strains have a strong bias for the *cmr*-ON orientation (∼70-96%, **Figure 7**). The chaining phenotype of Clade 5 strains prone to the *cmr*-ON state was enhanced during growth on a solid surface compared to broth culture (**Figures 4, 5**), consistent with prior analyses of the Clade 2 strain R20291, which exhibits greater *cmrRST* expression during growth on agar plates due to elevated c-di-GMP levels [46]. In addition, Clade 5 strains exhibit high levels of surface motility on agar plates, which is a *cmrRST-*induced phenotype in strain R20291 [33]. Finally, knocking-down the expression of the *cmrRST* operon in three Clade 5 strains biased to the *cmr*-ON state reduced their propensity to form chains (**Figures 7** & **S8**).

These findings lead to the question of what benefit *cmrRST* expression might confer to Clade 5 strains. In the Clade 2 R20291 strain, *cmrRST* expression is negatively correlated with flagellar motility, and growth conditions that favor flagellar motility select against the *cmr*-ON state [33]. Since Clade 5 strains lack flagellar motility [12], it is tempting to hypothesize that they are “primed” to form chains as a method to promote motility. Analyzing the regulation of c-di-GMP in different growth conditions in Clade 5 strains, particularly for TAL29996 relative to the other strains, will likely provide insight into the mechanisms that drive *cmrRST* expression, chaining, and surface motility in Clade 5 strains and the importance of these properties to these strains.

While the propensity of Clade 5 strains to form chains in a *cmrRST*-dependent manner might be expected to promote colonization or disease in mice based on studies of other Gram-positive pathogens [56, 57], we found that the frequency of *cmr-*ON cells decreases during the first 4 days of infection for strain TAL29600 (**Figure 8**). This suggests that there may be a selection against *cmr-*ON cells during the initial stages of murine colonization. Consistent with this hypothesis, a decrease in *cmr-*ON orientation was observed for Clade 2 strain R20291 during infection of hamsters [33]. However, since the frequency of *cmr-*ON TAL29600 cells increased in some mice at later stages of colonization (**Figure 8**), our data imply that the invertibility of the *cmrRST* switch region may promote *C. difficile*’s ability to adapt to different growth conditions. While it is possible that high levels of c-di-GMP during infection induce the expression of the *cmrRST* operon expression during murine infection [46], assessing whether Clade 5 strains form long chains during murine infection, for example, using fluorescence *in situ* hybridization [61, 62] or using transcriptional reporters to visualize the expression of the *cmrRST* operon at the single-cell level during infection would provide insight into these questions.

Indeed, the avirulence of most Clade 5 strains analyzed during murine infection (**Figure 8**) was somewhat surprising given that all of these strains were isolated from human patients experiencing CDI-related disease symptoms (**Table 1**). To our knowledge, very few Clade 5 strains have been analyzed during murine infection, with one study observing minor disease symptoms in mice for two Clade 5 strains three days post-infection, despite one of the strains causing severe disease in humans [20]. Interestingly, RT078 Clade 5 strains frequently cause asymptomatic infections in agricultural animals [32, 63] and mice may be important vectors of transmission in these settings [64, 65]. Thus, it is possible that Clade 5 strains are adapted for colonization rather than virulence in non-human systems. Consistent with this hypothesis, we found that the Clade 5 strains persist at high levels in the murine gut over time compared to the Clade 1 strain 630 (**Figure 8**, p < 0.005). Identifying factors that allow Clade 5 strains to grow more quickly would provide insight into whether their faster growth rate promotes their persistence in mice. It is also possible that increasing the inoculum could have allowed for a greater degree of disease severity to be observed and differences in virulence between the strains to be detected.

Interestingly, the chain length of Clade 5 strains did not correlate with their propensity to sporulate (**Figure 6**). The Clade 5 strains tested varied markedly in their sporulation frequencies, with most strains forming spores at frequencies >30% unlike strain TAL29600, which sporulates ∼5,000-fold less efficiently than the other Clade 5 strains (**Figure 6**). While little is known about the mechanisms regulating sporulation initiation outside of strains 630 and R20291, analyses of strain TAL29600, which sporulates poorly under laboratory conditions (**Figure 6**), could provide insight into the molecular determinants of sporulation initiation in Clade 5 strains. For example, differences in gene presence or polymorphisms in several c-di-GMP-related genes were observed in TAL29600 relative to the other Clade 5 strains (**Table S2**), and c-di-GMP has been implicated in regulating sporulation initiation events through unknown mechanisms [66, 67].

Our time-lapse microscopy analyses further revealed that a notable delay between cell division and cell separation is common to *C. difficile* strains, irrespective of their propensity to form long chains, because chains of two cells were frequently observed for Clades 1-4 and Clade 5 strain TAL29996 (**Figures S3** & **4**). Notably, these two-cell chains were typically segmented by DeLTA as a single cell because cell separation had not initiated, i.e. no invagination was detected (**Figure 4**, red dot). The loose coordination between cell division and cell separation in *C. difficile* relative to other bacteria likely relates to the absence of FtsEX homologs in *C. difficile*. In diverse bacteria, the FtsEX complex couples septal PG synthesis with PG hydrolases that mediate cell separation to result in a fast splitting of recently divided cells [68–71]. While it remains unclear whether coordination between cell division and cell separation exists in *C. difficile*, recent work has identified novel factors that control chaining in *C. difficile*. The CwlA peptidoglycan hydrolase mediates cell separation in *C. difficile* [72], and its export and thus activity is controlled by the Ser/Thr kinase PrkC [72]. The septum-localizing MldA or MldB proteins also promote chaining in *C. difficile* through unknown mechanisms [73], so future work could address whether the inefficient cell separation phenotype of Clade 5 strains is due to decreased CwlA export or MldA/MldB levels.

Importantly, these insights into the basic physiology of *C. difficile* were enabled by our development of a facile method for conducting time-lapse microscopy under anaerobic conditions. Since the growth chamber set-up involves commercially available GeneFrames and open-source software for conducting automated image analyses of time-lapse microscopy data [43, 44], the methods described in this manuscript can be applied to many anaerobic systems for studying the growth properties of diverse organisms and the impact of different growth conditions and mutant backgrounds on these properties. Our anaerobic set-up could be further coupled with recently developed, fluorogen-activated anaerobic imaging tags [74] to facilitate single-cell analyses of gene-specific transcription during anaerobic growth and dynamic protein localization studies [75]. Thus, there are many potential applications for the simple methods described here for studying the growth of anaerobes over time at the single-cell level.

## Materials and Methods

### Bacterial strains and growth conditions

All *C. difficile* strains were grown on brain heart infusion (BHIS) medium supplemented with 0.5% w/v yeast extract and 0.1% w/v L-cysteine with taurocholate (TCA; 0.1% w/v; 1.9 mM). Strains were sub-cultured into tryptone yeast extract (TY) broth supplemented with 0.1% w/v L-cysteine (TYC medium) prior to inoculation onto the time-lapse microscopy agarose pads. All strains were grown at 37°C under anaerobic conditions using a gas mixture of 85% hydrogen, 5% CO2, and 10% H2. For time-lapse experiments, 1.5% agarose pads supplemented with TYC medium were used as described above. Sporulation analyses were carried out on 70:30 medium (70% BHIS and 30% SMC) for 24h as described previously [76].

### Anaerobic time-lapse imaging of C. difficile growth

All imaging was carried out on a Leica DMi8 inverted microscope with a HC plan apochromat 63x 1.4 NA oil immersion phase contrast objective. Fluorescent membrane staining experiments were done with a Lumencor Spectra X light source, coupled with an XLED-QP quadruple-band dichroic beam-splitter (Leica) (transmission: 415, 470, 570, and 660 nm) along with an external emission filter wheel (Leica). FM4-64 was excited using a 470nm LED through a 470/20nm excitation filter and emitted light was filtered through a 590/50nm emission filter and captured with a Leica DFC9000GTC sCMOS camera. All experiments were carried out at 37°C using a microscope incubation system (Pecon), Leica Adaptive Focus Control hardware autofocus, and a high precision stage (Pecon) were used for all imaging experiments.

For time-lapse imaging of *C. difficile* growth, all bacterial strains were grown in 2 mL liquid TY medium to a turbid OD600 > 2-3; after 2 hours of growth, bacteria were diluted 1:100 for Clade 5 strains and all other strains were diluted 1:50 in fresh media and grown to mid-log phase (OD600 0.4-0.7).

An imaging chamber with a gas-tight seal was constructed using a 125 μL Gene Frame (Thermo Fisher) adhered to a glass slide generating a well for growth medium. The slide was then transferred to the anaerobic chamber. In the anaerobic chamber, the gene frame was filled with 500μl 1.5% Top vision low melting point agarose and tryptone yeast extract media containing 0.1% w/v L-cysteine to scavenge oxygen and maintain anaerobic conditions. While the agarose was molten, a second clean slide was placed over the top and the agar pad was placed on a frozen small freezer block (for holding PCR strip tubes) for 10-30 minutes until the agarose-media mixture was solid. For experiments using FM4-64, agarose pads were made the same way, with the addition of FM4-64 to a final concentration of 1 μg/mL directly to the agarose/media solution prior to making the agar pad.

The agar pad was dried for 5-10 minutes until visible liquid on the surface of the pad was evaporated. 1 μL of mid-log cells were spotted on the pad, dried, and a #1.5 coverslip (VWR) was adhered to the Gene Frame. The cells were imaged at 37°C until they reached confluency in the field of view. This was anywhere from 2.5 hours for Clade 5 strains to 6 hours for Clades 1-4 for all experiments with images taken at 5-minute intervals.

### Image analysis, computing hardware, and statistical analysis

All movie frames were trimmed to the point when cells were not overlapping and out of focus regions were cropped. The resulting images were analyzed using the Python library DeLTA 2.0 [43, 44]. All image and data analyses were done on a PC running Windows 10 equipped with an AMD Ryzen 5900HX 8-core CPU, 32GB DDR4 RAM, 2 1TB NVME SSDs, and an NVIDIA RTX3080 GPU with 16GB VRAM. Analysis of the output data and data visualizations were done in Python using Matplotlib/Seaborn, Pandas, Numpy, Scipy, and the Statannotations library.

### Septum detection during live-cell microscopy

The image processing starts with the masks generated using DeLTA. First, we performed erosion on the mask with a disk of radius of 1 pixel to avoid effects of membrane fluorescence. The photo was cropped following the eroded mask contour. Hence, only the interior of the contour was considered. Pixel values were rescaled such that the minimum pixel value of inner pixels was mapped to 0 and the maximum was mapped to 255. We set a threshold intensity as the pixel value of the 95% quantile of pixel intensities. The image was slightly blurred by convolution with a Gaussian filter with standard deviation of 2 pixels. The septum corresponds to the contours found after performing an adaptive thresholding based on thresholds defined using a Gaussian-weighted method. This thresholding was performed using the open cv2 library in Python. The resultant contours were manually verified from videos generated using the tracking from the outputs from DeLTA. The manual validation mainly involved filling in the time points where the algorithm missed a ring that was visible in the video. We did not assume the existence of a ring if the algorithm did not detect it in a previous frame.

### Cell length estimation

We measured the cell projected area from the DeLTA contours of the images as the pixel amount of the contours. However, estimating cell length was challenging because some cells were very long and bent. To overcome this problem, we selected 30 images of three different cells (from strains 630, TAL3050 and TAL28131) that were straight and had different lengths. For these cells, we calculated the cell length as the longest side of the minimum bounding rectangle of the contour. From these lengths of straight cells, we also estimated the best cell width as the average of the projected area divided by the length. Considering the extreme cell length, the effects of the rounded tips were negligible and the rectangle shape adequately approximated length. We verified that this mean value showed low variability for the three strains. Then, we used this width value to estimate the length of all the cells, including the bent ones, by dividing their projected area by the width. This way, we obtained a consistent measure of cell length that was independent of bending.

### Elongation rate estimation

We tracked cell size over time and identified the division points as the ones where the cell size (projected area) dropped by more than 30% compared to the current cell size value. We fitted an exponential function (with base e) of time to the data points between two divisions and estimated the elongation rate from the exponent of the best fit. We expressed the elongation rate in doublings/hr, which means how many times the cell size doubles in one hour. For example, an elongation rate of 2 doubling/hr means that the cell size doubles two times in 1 hr, which corresponds to an exponent of 2ln(2) 1/hr. For the statistics, we only included the elongation rates that had a high quality of fit, with an R2 coefficient greater than 0.9.

### Bulk growth measurements

Starter cultures were grown until early stationary phase in BHIS (or TYC medium as indicated) then diluted 1:50 into BHIS (or TYC medium). For the CDDM growth analyses, starter cultures were prepared in CDDM medium at a relatively high density and then back-diluted 1:25 into CDDM. When the cultures (for all three media conditions) reached an OD_600_ of 0.5, they were diluted 1:50 into 200 µL of either BHIS, TYC, or CDDM in a flat 96 well polystyrene plate (CellTreat). The OD_600_ was analyzed every 15 min for 24 hrs in a BioTek Epoch plate reader with shaking. Bulk growth measurements are based on a minimum of three independent replicates across a minimum of 2 experiments. The growth rate was calculated from the linear range of the growth curves, between 105 min to 180 min of growth.

### Cell wall labeling

HADA (Tocris Bioscience) was added to exponentially growing cell culture to a final concentration of 50-100 µM and incubated for ∼2 mins before cell fixation. Cells were fixed as previously described [77]. Briefly, 500 µL of cell suspension was added to 120 μL of a 5X fixation solution containing paraformaldehyde and NaPO_4_ buffer. Samples were mixed and incubated in the dark for 30 min at room temperature, followed by 30 min on ice. Fixed cells were washed three times in phosphate-buffered saline (PBS) and resuspended in ∼50 µL of PBS. Cells were imaged within 72 hours after fixation.

### Sporulation assays

Starter cultures were grown until early stationary phase in BHIS then diluted 1:50 into BHIS. When the cultures reached an OD_600_ between 0.35 and 0.75, 120 µL of the culture was spread onto 70:30 (70% SMC media and 30% BHIS media) agar plates (40 ml media per plate) and then incubated for 20-24 hrs before the sporulating cells were scraped from the plate into phosphate-buffered saline (PBS). Sporulation levels were visualized by phase-contrast microscopy as previously described [78].

### Heat resistance assay

Heat-resistant spore formation was measured 20-24 hrs after sporulation was induced on 70:30 agar plates as previously described [76]. The percent sporulation of given culture represents the ratio of heat-resistant colony-forming units (CFUs) to total CFUs. Percent sporulation was determined from a minimum of 3 biological replicates.

### Spore purification

Spores were purified as previously described [79] by scraping up sporulating cells incubated on 70:30 medium for 3 days into ice-cold H_2_O. The cells were washed several times in ice-water over the course of a day and incubated on ice overnight. The following morning, the sample was pelleted, and cells were suspended in 1 X DNAse buffer (New England Biolabs) and then treated with DNAse (New England Biolabs) for 30 min at 37°C. The samples were washed one more time before being resuspended in 20% Histodenz and then layered onto a 50% Histodenz layer. The resulting mixture was pelleted, and the supernatant was aspirated off using a vacuum aspirator. The pelleted spores were washed in ice-cold water 2-3 times and the optical density of the purified spores was measured.

### Genomic DNA preparation

Starter cultures were grown until early stationary phase in BHIS then back-diluted 1:50 into BHIS and grown until an OD_600_ of around 0.7-0.8 was reached. 10 mL of the culture was pelleted and then frozen at −80°C. After thawing the sample, it was resuspended in a 25% sucrose TE buffer (10 mM Tris, 1mM EDTA), incubated with 100 mg/mL lysozyme for 37°C for 1 hr. After the cultures tarted to lyse, proteinase K, RNAse A, EDTA, Sarkosyl, and NaCl was added. Phenol:Chloroform:IAA (25:24:1) was added to extract proteins, gently mixed, and then the sample was pelleted to separate the phenol and aqueous layer. The aqueous layer was then added to Chloroform:IAA (24:1), mixed gently, then centrifuged. The aqueous layer was then precipitated using isopropanol and incubated at −20°C for a minimum of 15 min. The precipitated DNA was pelleted and then washed with 70% ethanol. The pellet was air dried and then gently resuspended in 10 mM Tris pH 8.0 elution buffer.

### Genomic analyses

Genomic DNA was sequenced by MiGS at the University of Pittsburgh (now SeqCenter) according to their standard protocol. Libraries were sequenced on an Illumina NextSeq 500 platform to generate paired-end 150 bp reads. Illumina reads of RT078 genomes were assembled into contigs using SPAdes (v3.13.0),[80] and genes were called and annotated using Prokka (v1.11) [81]. Assembled and annotated contigs of five RT078 strains (TAL28131, TAL29600, TAL29996, TAL30550, TAL30574) were applied for pangenomic analysis. Default settings were used based on the Anvi’o workflow for microbial pangenomcis with adjustments for minbit as 0.5 and mcl-inflation as 10[82–84]. For SNPs analyses, reads of five RT078 genomes were aligned to the reference M120 and variants were called by breseq (v. 0.38.1) by default settings.[85]

### qPCR analyses

Each genomic DNA sample was analyzed by qPCR with primers that amplify the *cmr*-ON sequence orientation, the *cmr*-OFF orientation, or the reference gene *rpoA* [33]. Each 20-µL qPCR reaction consisted of 100 ng genomic DNA, 100 nM primers, and SensiMix™ SYBR reagents (Bioline). The reactions were run on a LightCycler® 96 (Roche Diagnostics), and *cmr* switch orientation frequencies were calculated as described previously [46].

### Surface motility assays

Starter cultures were grown until early stationary phase in BHIS then back-diluted 1:50 into BHIS and grown until an OD_600_ of 0.5 was reached. 10 µL of the exponential-phase cultures were then spotted onto BHIS plates and incubated at 37°C for 5 days after which the plates were scanned using a flatbed scanner.

### Mouse infection experiments

Mouse experiments were performed under the guidance of veterinary staff within the Tufts Comparative Medicine Services (TCMS) core. All animal studies were done with prior approval from the Tufts Institutional Animal Care and Use Committee (IACUC protocol #B2024-30). Conventional 7-week-old C57BL/6 female mice from Jackson Laboratories were housed in a sterile (autoclaved cage and bedding) large cage (24”x17”) with autoclaved water and irradiated food (Teklad 2918) for 10 days to allow for normalization of microbiota across mice through coprophagy. After the 10-day normalization period, mice were started on cefoperazone, which was added to their water at a concentration of 0.5mg/ml. Mice were allowed to drink the cefoperazone water ad libitum for 10 days, after which they were placed back on sterile water without antibiotic. After a 2-day period of being on normal sterile water, mice were weighed and given a single dose of clindamycin (10mg/kg) via intraperitoneal injection. Immediately after IP injection, mice were moved to standard-size autoclaved mouse cages (4 mice per cage) with sterile food and water. 24 hours following the clindamycin injection, mice were inoculated with 1 x 10^5^ spores of *C. difficile* (in 1xPBS) via oral gavage using a metal, reusable needle. Mice were weighed by being placed in a plastic Nalgene cup on top of a scale. Fecal pellets were collected just prior to oral gavage to ensure no prior *C. difficile* colonization and to note a baseline weight. Following inoculation, mice were weighed to monitor % weight change over time on days 1-4, 7, and 14. Fecal pellets were collected in duplicate on the same days for *C. difficile* CFU enumeration and qPCR for detection of *cmr* switch orientation. On day 14, a terminal weight was taken, and a fecal pellet was collected, followed by sacrifice via CO_2_ inhalation with cervical dislocation as secondary method of euthanasia. Two experimental replicates were completed using 4 mice per group, resulting in a total of 8 mice per condition.

### C. difficile CFU enumeration from mouse fecal pellets

*C. difficile* engraftment in mice was monitored over a 14-day period. Fecal pellets were collected from mice on days 0-4, 7, and 14. Each pellet was weighed and then suspended in 1x PBS. 10 µL of the suspension was then serially diluted 1:10 in 1x PBS in a 96 well plate, and 5 µL was spotted onto TCCFA agar to select for *C. difficile*, such that the dilutions plated were 2 x 10^3^ – 2 x 10^7^. *C. difficile* colonies were counted 24 hours after plating to allow for sufficient growth. *C. difficile* CFUs were normalized by gram of fecal material.

### Data accessibility statement

The data set and data processing scripts are publicly available at https://doi.org/10.5281/zenodo.13352469 [86]. Genomes are available from NCBI under the BioProject ID PRJNA1238863.

## Acknowledgements

We are grateful to Dr. Cheleste Thorpe, Dr. Eric Pamer, Dr. Bruno Dupuy, Dr. Lynn Bry, and Dr. Trevor Lawley for providing the strains used in this study.

