## Supplemental Figures for "Unique growth and morphology properties of Clade 5 *Clostridioides difficile* strains revealed by single-cell time-lapse microscopy"

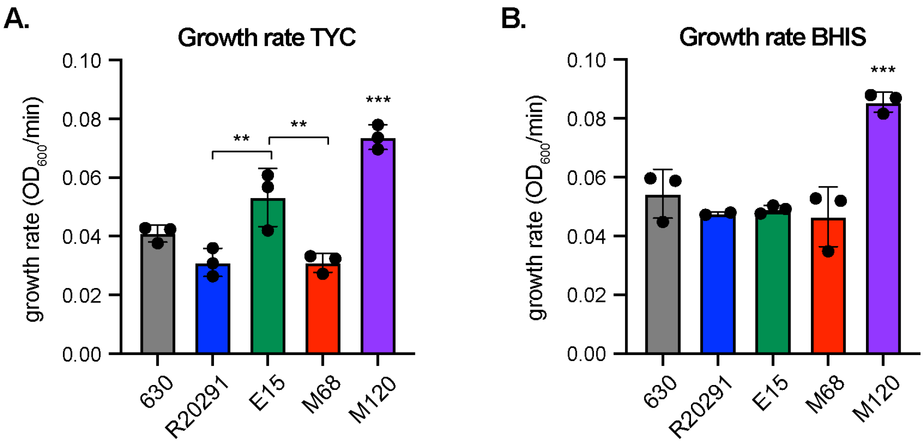


**Figure S1. Growth rates measured from optical density-based analyses of bulk population growth in TYC and BHIS media.** The growth rate was calculated from the linear range of the growth curves.

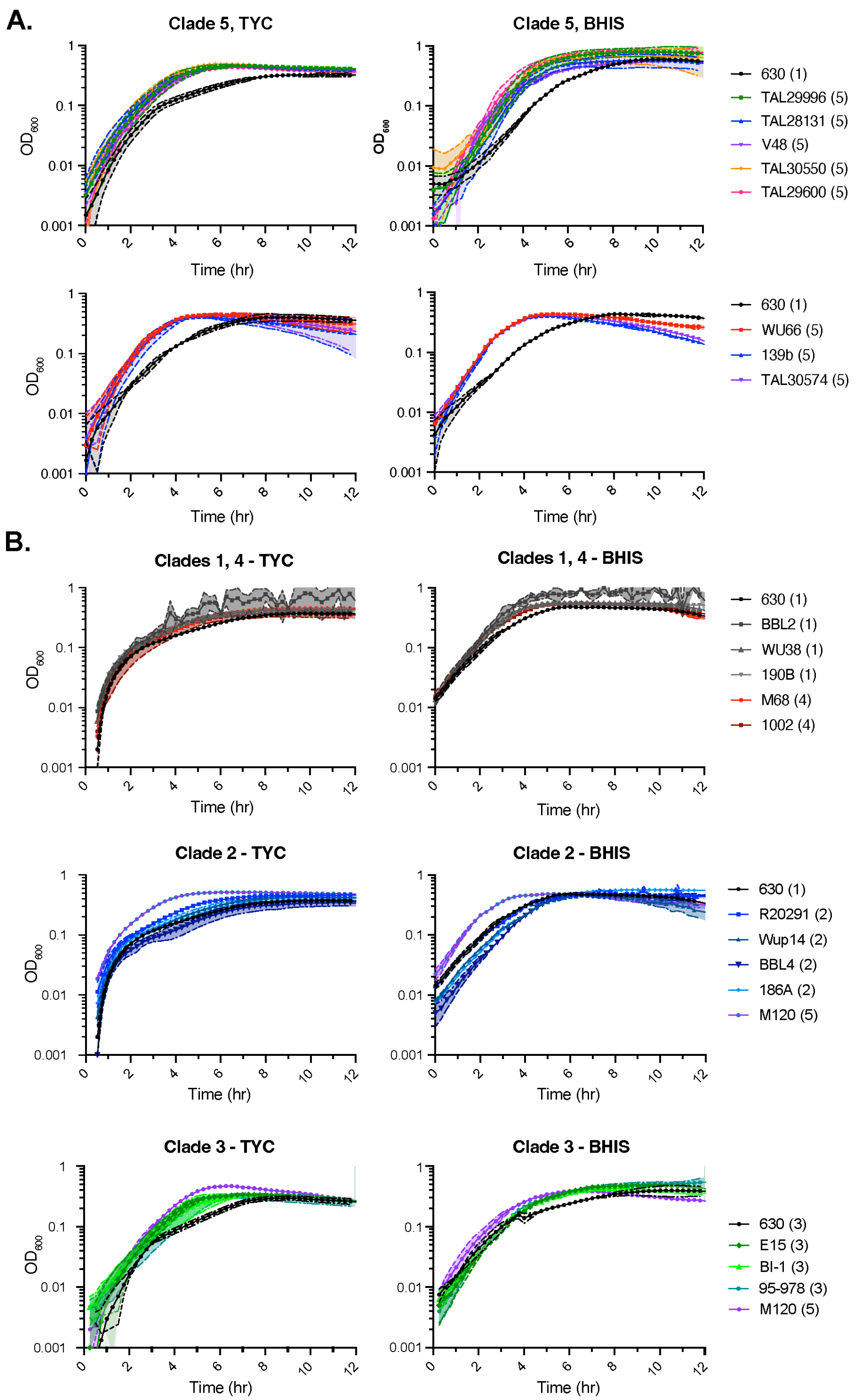


**Figure S2. Optical density-based analyses of bulk population growth of the indicated strains during growth in TYC and BHIS media**. The number in brackets indicates the clade to which a given strain belongs. For the Clade 3 growth curves, the data shown for Clade 1 strain 630, Clade 3 strain E15, and Clade 5 strain M120 is identical to the data shown in Figure 2.

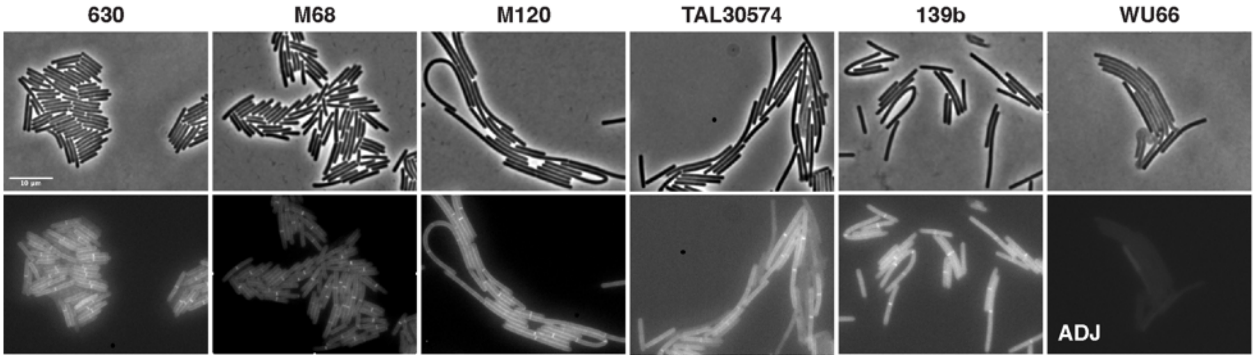


**Figure S3. Clade 5, but not Clade 1 or 4, strains form chains during logarithmic growth in BHIS broth culture**. Representative micrographs showing phase-contrast (top) and peptidoglycan labeling with the fluorescent D-amino acid, HADA (bottom) following growth in rich medium (BHIS) to logarithmic phase. Scale bar, 10 µm. Data are representative of multiple independent experiments. **ADJ** indicates that the brightness of the image was enhanced to detect HADA labeling in WU66.

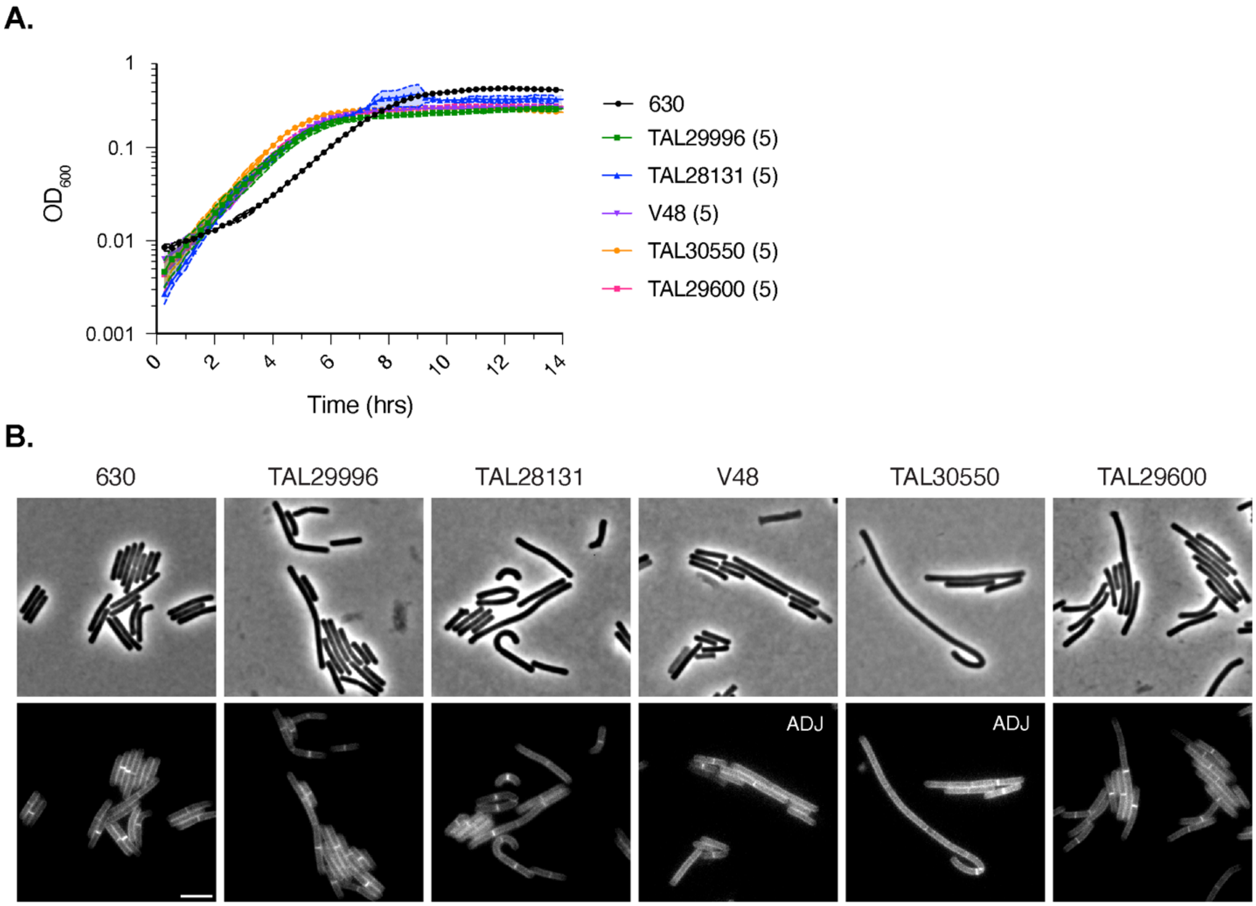


**Figure S4. Clade 5 strains form chains even when growth is slowed during growth in minimal medium broth culture.** (A) Optical density-based analyses of bulk population growth of the indicated strains during growth in CDDM minimal medium. (B) Representative micrographs showing phase-contrast (top) and peptidoglycan labeling with the fluorescent D-amino acid, HADA, (bottom) following growth in CDDM to logarithmic phase. All strains shown, with the exception of the Clade 1 strain 630, are Clade 5 strains. Scale bar, 5 µm. Data are representative of three independent experiments. **ADJ** indicates that the brightness of the image was enhanced to detect HADA labeling in V48.

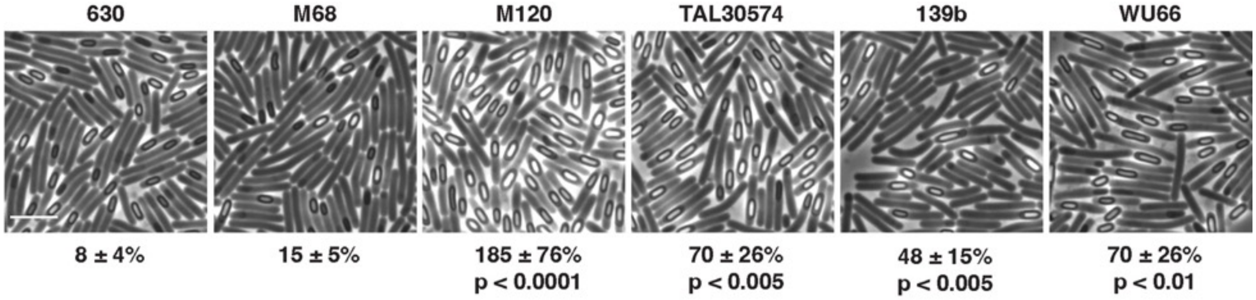


**Figure S5. Sporulation levels in clinical isolates grown on 70:30 medium**. Phase-contrast microscopy of the indicated strains ~24 hrs after sporulation induction. The percent heat-resistant spores is indicated below the respective images. The percentage was determined from 20-24 hr sporulating cultures and represent the mean and standard deviation for a given strain based on a minimum of three biological replicates. Statistical significance relative to Clade 1 strain 630 was determined using a one-way ANOVA and Tukey's test. M68 is a Clade 4 strain. The remaining strains are Clade 5 strains. It should be noted that the image shown here for Clade 1 strain 630 is the same as in Figure 6. The scale bar represents 5 µm.

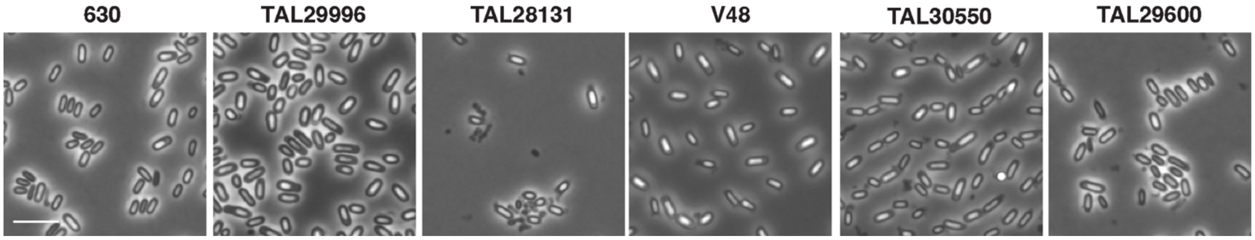


**Figure S6. Spores purified from Clade 5 strains.** Spores were purified using a Histodenz gradient. Strain 630 is provided as a reference. The scale bar represents 5 µm.

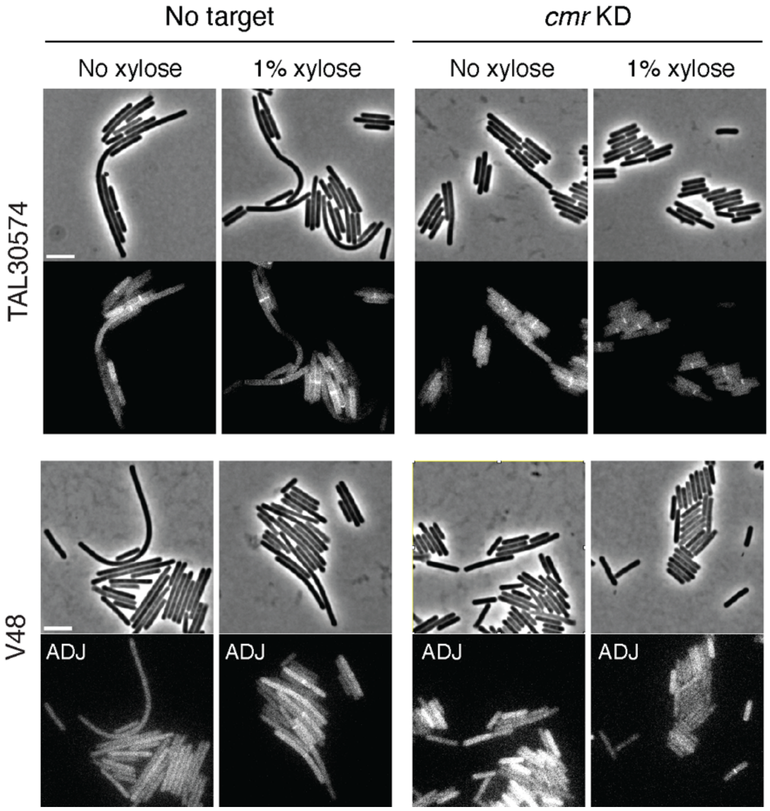


**Figure S7. Expression of the *cmrRST* operon promotes cell chaining.** Representative micrographs of *cmrRST* CRISPRi knock-down strains in the indicated Clade 5 strain background compared to a no target control. Phase-contrast (top) and peptidoglycan labeling with the fluorescent D-amino acid, HADA, (bottom) images following growth in rich broth (BHIS) to mid-logarithmic phase. Scale bar, 5 µm.

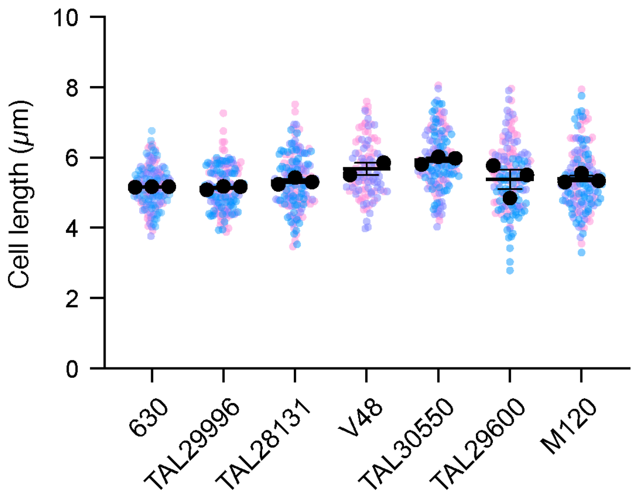


**Figure S8. Cell length during growth on 70:30 sporulation medium**. The cell length of a minimum of 50 cells per replicate was measured for the indicated strains after 20-24 hrs of growth on 70:30 sporulation medium. Black circles indicate the mean cell length for a given replicate is plotted with black circles; black lines indicate average means.

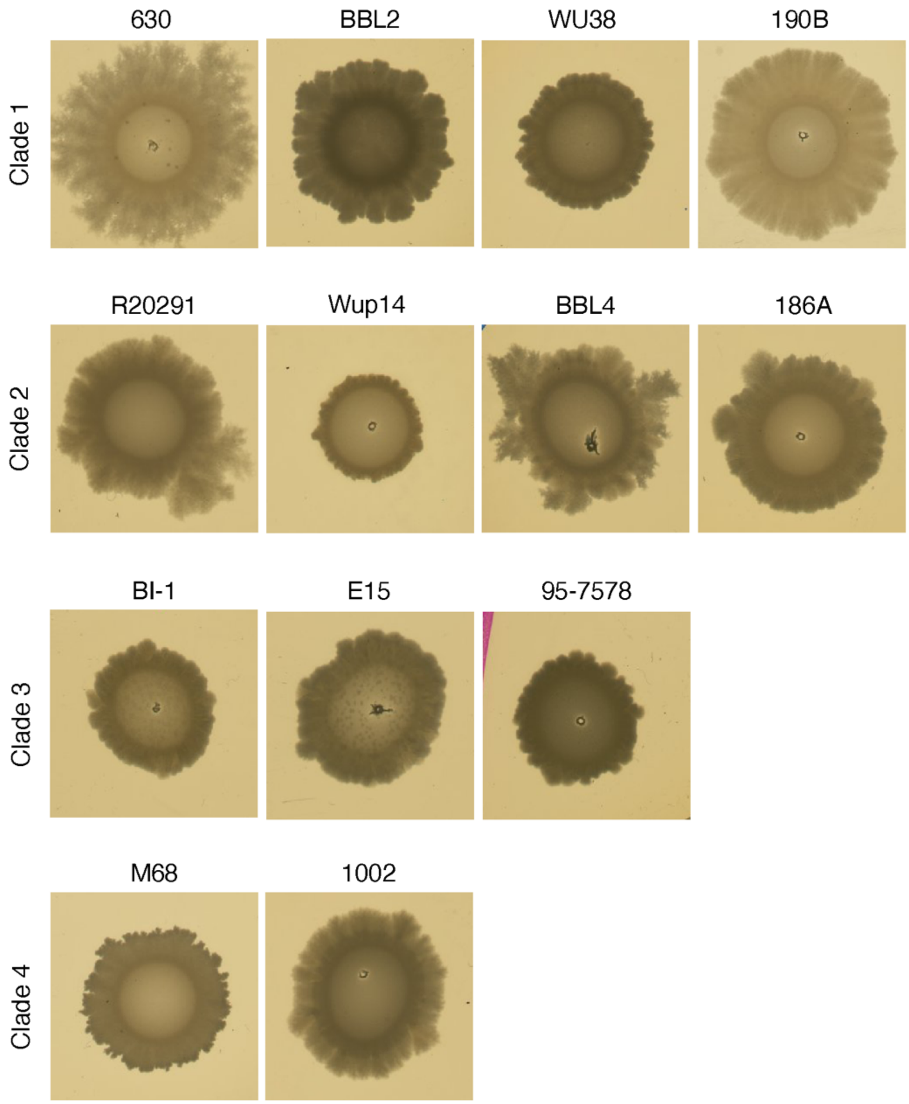


**Figure S9. Surface motility in Clade 1-4 strains.** Representative images of surface motility 5 days after exponentially growing liquid cultures were spotted onto BHIS agar plates.

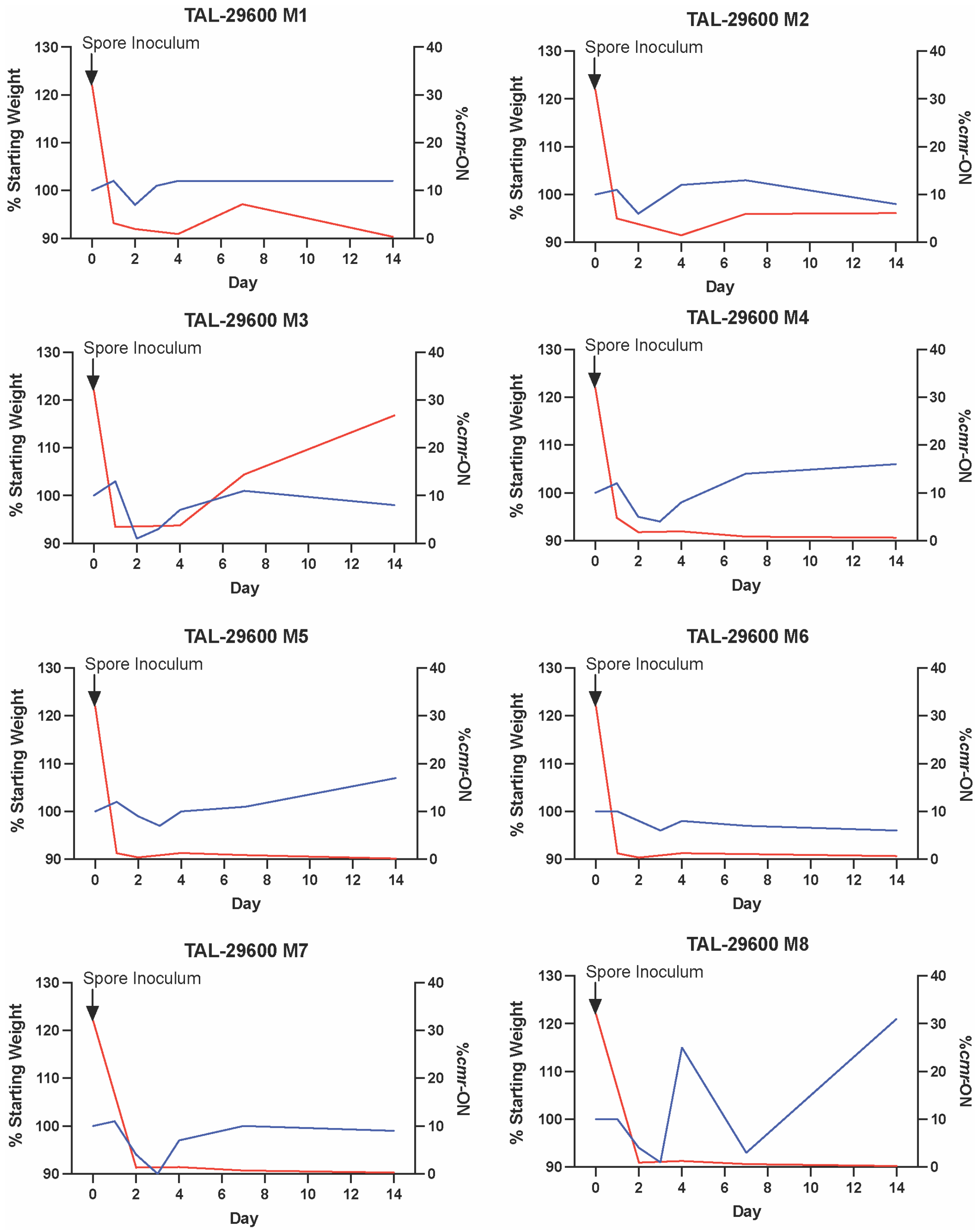


**Figure S10. Infection and *cmr* switch orientation dynamics during TAL29600 infection.** Percent weight loss and *cmr* ON switch orientation measured in fecal pellets of individual mice infected with TAL29600 on the indicated days by qPCR. For the strain TAL29600 inoculum, 32% of the spores had the *cmr* ON spores

**Table S1. Average nucleotide identity for orthologous genes for Clade 5 strains**

| | TAL28131 | TAL29600 | TAL29996 | TAL30550 | TAL30574 |
|---|---|---|---|---|---|
| TAL28131 | 100% | 99.83% | 99.96% | 99.94% | 99.88% |
| TAL29600 | 99.90% | 100% | 99.995% | 100.00% | 99.92% |
| TAL29996 | 99.95% | 99.996% | 100% | 99.99% | 99.99% |
| TAL30550 | 99.96% | 99.99% | 99.98% | 100% | 99.96% |
| TAL30574 | 99.85% | 99.92% | 99.99% | 99.98% | 100% |

**Table S2. Breseq analyses of Clade 5 strain genomes**
The M120 genome was used as the reference for these genomic comparisons.

**Table S3. Anvi'o analysis of the accessory genome identified for the Clade 5 strains sequenced**.
